# Adaptive strategies of Caribbean sponge holobionts beyond the mesophotic zone

**DOI:** 10.1101/2025.01.31.635867

**Authors:** Benoît Paix, Alexane Thivet, Celso Domingos, Özlem Erol, Niels van der Windt, Young H. Choi, Nicole J. de Voogd

## Abstract

Marine sponges and their microbiomes function together as holobionts, playing essential roles in ecosystem dynamics and exhibiting remarkable adaptability across depth gradients. This study utilized a multi-omics approach, integrating microbiome and metabolome analyses, to investigate adaptive strategies in sponge holobionts inhabiting the mesophotic (80-125 m), upper-rariphotic (125-200 m), and lower-rariphotic (200-305 m) zones of Curaçao. We hypothesized that depth-related environmental factors drive distinct adaptive strategies, similar to patterns observed in fish and coral assemblages.

Results revealed major differences in holometabolomes and microbial communities between Demospongiae and Hexactinellida sponges, reflecting class-specific adaptive strategies. Notably, phospholipid homeoviscous adaptation to temperature and pressure emerged as a key mechanism in phosphorus metabolism. Adaptations in nitrogen metabolism were linked to diverse ammonia oxidizing archaea (AOA) symbionts, and dissolved organic matter cycling. Hexactinellid microbiomes exhibited intra-specific heterogeneity; however, species-specific associations with AOA symbionts such as *Cenarchaeum* and *Nitrosopumilus* were observed. Additionally, the lower-rariphotic hexactinellid holometabolomes highlighted the significance of the nested ecosystem concept through the identification of secondary metabolites produced by their associated fauna (aphrocallistins by zoanthids, and xanthurenic acid by shrimp).

This study highlights the ecological significance of sponge holobionts in mesophotic and rariphotic ecosystems, revealing diverse adaptations to unique physicochemical conditions and biotic interactions.

## Introduction

Marine sponges (Phylum Porifera) are recognized as vital ecosystem engineers in benthic environments, providing essential ecological functions that support diverse marine life. Their highly efficient filter-feeding activity significantly contributes to the cycling of dissolved organic matter (DOM) and nutrient transfer within marine ecosystems (de Goeij et al., 2013; Pawlik & McMurray, 2020). Sponges host a rich diversity of macrobenthic fauna (Goren et al., 2021; Ilan et al, 1999, Kandler et al 2019) and complex microbial communities (Thomas et al., 2016; Webster & Thomas, 2016). Their associations with symbiotic microbes form a tightly integrated unit known as the holobiont (Rosenberg & Zilber-Rosenberg, 2018; Zilber-Rosenberg & Rosenberg, 2008; 2021). Microbial symbionts play a crucial role by producing essential compounds such as vitamins (Wilkinson & Fay, 1979; Fan et al., 2012) and diverse secondary metabolites that function as chemical defenses against predators, competitors, and biofouling organisms (Hardoim & Costa, 2014; Pita et al., 2018). This functional diversity enables sponges to adapt to challenging environments, particularly under rising ocean temperatures (Bell et al., 2013) and ocean acidification (Botté, 2019; Posadas et al., 2022). Moreover, sponge microbiomes facilitate the assimilation of specialized organic compounds, enabling survival in nutrient-poor environments (Rix et al., 2017; Bart et al., 2021).

While these adaptations are relatively well-studied in shallow-water ecosystems, research on sponge microbiomes in mesophotic and deep-sea environments remains limited (Steinert et al., 2020; Garritano et al., 2023; Dies-Vives & Riesgo 2024). Expanding this knowledge is crucial to fully understand sponge adaptation and resilience in response to changing ocean conditions. In the mesophotic zone (30–150 m), reduced light levels lead to shifts in sponge composition compared to shallow water sponges. While Demospongiae remain dominant in mesophotic sponge communities, their association with photosynthetic microbes declines. Instead, they rely more on heterotrophic and chemosynthetic symbionts, such as Chloroflexi and Acidobacteria, for nutrient acquisition and organic matter cycling (Moitinho-Silva et al., 2017; Cleary et al., 2023). Beyond 200 m, glass sponges (class Hexactinellida) become more prevalent. Adapted to cold, high-pressure, and nutrient-limited conditions of deep-sea environments, these sponges form symbioses with specialized microbial taxa, such as ammonia-oxidizing archaea (AOA) and ammonia-oxidizing bacteria (AOB), which initiate nitrification by converting ammonium (NH₄⁺) into nitrite (NO₂^-^) (Steinert et al., 2020; Tian et al., 2016; Garritano et al., 2023). Despite their ecological significance, the microbiomes of Hexactinellida remain poorly understood and exhibit a more heterogeneous microbial distribution than other Low Microbial Abundance (LMA) sponges (Busch et al., 2022; Tian et al., 2016; Bayer et al., 2020; Steinert et al., 2020).

The rariphotic zone (150–300 m), first described in the Caribbean Sea by Baldwin et al. (2018), serves as a transitional layer of reef-fish communities between the mesopelagic zone (twilight zone) and the bathypelagic zone (midnight zone). This zone refers to a specific depth range characterized by low irradiance levels, where light penetration becomes insufficient to support photosynthesis (Baldwin et al., 2018). Recent observations also suggest a broadening of this delineation to other benthic communities on a wider scale in the Caribbean (Semmler et al., 2017; Stefanoudis et al. 2019). However, it remains unclear how the rariphotic zone can influence other benthic communities beyond reef fishes. Historically, Caribbean sponge diversity has been primarily documented in shallow coral reef ecosystems (van Soest, 1981), leaving the deep Caribbean reef slopes relatively unexplored. However, recent advancements in deep-sea exploration have led to the discovery of new species, including glass sponges emerging first in the rariphotic zone (van Soest, Meesters, & Becking, 2014; Reiswig & Dohrmann, 2014; Cleary et al., 2024a). In particular, the community structure and functions of sponge microbiomes are poorly understood in rariphotic depths. Investigating these relationships could reveal novel insights into adaptive strategies, symbiotic interactions, and the production of bioactive compounds, especially under the unique environmental pressures of this transitional depth zone (Hardoim & Costa, 2014; Steinert et al., 2020).

To explore adaptive strategies, this study employed a multi-omics approach integrating microbiome and metabolome analyses to investigate sponge holobionts in mesophotic and rariphotic depths. The reefs of Curaçao were selected as an ideal study site due to (i) the mesophotic-rariphotic delineation previously described in this area (Baldwin et al., 2018), and (ii) the coexistence of diverse Demospongiae and Hexactinellida communities, providing a unique opportunity to compare phylum-specific adaptive strategies (van Soest et al., 2014). This study aims to assess whether the mesophotic-rariphotic boundary, previously linked to low connectivity in fish and coral reef communities (Baldwin et al., 2018; Semmler et al., 2017; Stefanoudis et al. 2019), also applies to sponges. We hypothesize that sponge classes exhibit distinct adaptive strategies below the mesophotic zone. These strategies could be driven by key metabolic pathways shaped by microbiome interactions in response to environmental gradients such as temperature, pressure, nutrient availability, oxygen concentration, and light limitation.

Sponge samples were collected from three defined photic zones: (i) Mesophotic zone (80-125 m), (ii) Upper-rariphotic zone (125-200 m), and (iii) Lower-rariphotic zone (200-305 m) (Baldwin et al., 2018). Sponge samples were analyzed using a multi-omics workflow that combined 16S rRNA gene metabarcoding and UHPLC-HRMS metabolomics (**Figure S1**), enabling a comprehensive assessment of microbial and chemical diversity to uncover sponge adaptive mechanisms across depth gradients.

## Material and methods

### Sample collection and processing

A total of 60 sponge specimens were collected across four dives (**Table S1**): the first in October 2018 (n=2), two in April 2022 (n=10 and 34), and the final dive in March 2023 (n=14). Sampling covered an area of approximately 0.5 km^2^ off the Curaçao substation (12.083197 N, 68.899058 W, **Figure S2**). Collections were conducted with the Curasub, a submersible equipped with two robotic arms for collecting sponge specimens, along with a compartmentalized basket and storage container for safe transport during dives (http://www.substation-curacao.com/). Sponges were collected from depths of 89 to 305 meters, the latter being the maximum depth accessible by the Curasub. Specimens were initially identified by phylum during collection, resulting in 24 Demospongiae and 36 Hexactinellida samples. Specifically, 12, 5, and 7 demosponge specimens were collected from the mesophotic, upper- and lower-rariphotic zones, respectively. For Hexactinellida, 11 and 25 specimens were collected from the upper- and lower-rariphotic, respectively. Due to the logistical challenges of this sampling type and the patchy distribution, not all species could be collected in triplicate (Robertson et al., 2022; **Table S1**). Sponges were photographed both during collection and after retrieval to the surface (**Figure S3**). Tissues were subsampled for morphological, molecular, and metabolomics analyses using sterilized tweezers and scalpel blades. Samples were immediately preserved in 5 ml tubes containing 96% ethanol and stored at -20 °C until further processing. Extraction blanks for molecular and metabolomics analyses were prepared without sponge tissues. Additional samples of sponge-associated fauna (shrimps and zoanthids) were also photographed and preserved using the same protocol as the sponge specimens (**Figure S4**).

### DNA extractions

Sponge tissues were sectioned into pieces of approximately 3*2*1 mm, using sterilized tweezers and scalpel blades, yielding an average wet weight of 0.14 g (SD: ±0.02). Microbial and sponge DNA were extracted using the DNA^TM^ SPIN Kit for Soil (MP Biomedicals, Inc.) following the manufacturer’s instructions. The same protocol was applied to two DNA extraction blanks to monitor contamination. Additionally, DNA from Hexactinellida specimens was specifically extracted for barcoding identification using the DNeasy (Qiagen) Blood and Tissue Kit. All DNA extracts were stored at -20 °C until further processing.

### Sponge barcoding and morphological identification

Sponges were identified based on external morphology, spicules, and skeleton preparation following standard procedures and for Hexactinellida outlined by Reiswig & Stone (2013), following identification keys from *Systema Porifera* (Reiswig, 2002[2004]) and checking all species with the World Porifera database (de Voogd et al., 2025).

For the barcoding of demosponges and sponge-associated fauna, the 28S rRNA gene was amplified using the primers 28S-C2-fwd and 28S-D2-rev (Chombard et al. 1998), following the protocol described by Maslin, Paix et al., 2024 (initial denaturation: 98℃ for 30 s, 30 cycles of denaturation at 98℃ for 10 s, 51℃ for 10 s, 72℃ for 15 s, and final extension of 72℃ for 5 min). PCR1 products were checked using E-Gel^TM^ (agarose gels at 2%), and the absence of amplicons was validated for the negative controls and two extraction blanks. PCR1 products were cleaned using magnetic beads on the C.WASH plate washer (CYTENA, GmbH). PCR2 were performed with an initial denaturation step of 3 min at 95°C followed by 15 cycles of 15 s at 95°C, 15 s at 62°C and 50 s at 65°C, and a final extension step of 3 min at 65°C. Samples from each plate were pooled by collecting 1 µl per sample using the OT-2 Liquid Handler (Opentrons Labworks, Inc.) and cleaned using NucleoMag NGSBeads. End-repair and dA-tailing were performed on the pools using the NEBNext Ultra II End Prep Kit, followed by another clean-up. The DNA concentration of the pooled samples was quantified using a TapeStation 4150 (Kit HSD 5000, Agilent Technologies, Santa Clara, CA, United States) in combination with the Qubit dsDNA BR assay kit (Thermo Fisher Scientific, USA) to take forward an equimolar mass to the native barcode ligation step, performed using the NEB Blunt/TA Ligase Master Mix. Pools were then combined, followed by an AMPure XP bead clean-up (0.4x ratio). Lastly, native adaptors for sequencing were ligated using the NEBNext Quick ligation kit, and a final quantification of the library was performed on the Qubit Flex. The final library size was set up to 12 μl at 10-20 fmol for sequencing on a Nanopore R10.4.1 flow cell. Consensus sequences were generated by assembling raw reads using NGSpeciesID 0.3.0 and medaka 1.8.0.

For hexactinellids, PCR amplifications were conducted following the protocol of Dohrmann et al., (2006) using the primers 16S1fw and 16SH_mod for the amplification of the 16S rRNA gene (Bridge et al., 1992; Dohrmann et al., 2008). The resulting PCR products were sequenced *via* Sanger sequencing by STABVIDA using the BigDye Terminator v 3.1 kit on an ABI 3730XL DNA analyser (Applied BiosystemsTM, Foster City, CA, USA).

The sequences generated were analyzed using the Basic Local Alignment Search Tool (BLAST) (https://www.ncbi.nlm.nih.gov/genbank/) to assess similarity with existing sequences in public databases and confirm the taxonomic placement of the species.

### Library preparation and high throughput sequencing for 16S rRNA gene metabarcoding

The library preparation for metabarcoding analyses was conducted through a two-step PCR protocol for all samples, two extraction blanks, and a negative control. For PCR1, the V4-V5 regions of the bacterial and archaeal 16S rRNA gene were targeted with the 515F-Y/926R primers (Parada et al. 2016). PCR1 reactions were performed with the KAPA HiFi HotStart Ready Mix PCR Kit (Roche Molecular Systems, Inc.) in a T100 Thermal Cycler (Bio-Rad, Hercules, CA, United States). The following thermal cycling scheme was set up: initial denaturation at 95°C for 3 min, 30 cycles of denaturation at 98°C for 20 s, annealing at 50°C for 30 s, followed by extension at 72°C for 30 s. The final extension was carried out at 72°C for 5 min. PCR1 products were checked using E-Gel^TM^ (agarose gels at 2%), and the absence of amplification was validated for the negative controls and two extraction blanks. PCR1 products were then cleaned using NucleoMag NGS-Beads (bead volume at 0.9 times the total volume of the sample, Macherey Nagel, Düren, Germany) and the VP 407AM-N 96 Pin Magnetic Bead Extractor stamp (V&P Scientific, San Diego, CA, United States). PCR2 were performed using IDT xGen^TM^ NGS Adapters & Indexing Primers kit (Integrated DNA Technologies, Inc.) with an initial denaturation step of 3 minutes at 95°C followed by 8 cycles of 20 seconds at 98°C, 30 seconds at 55°C and 30 seconds at 72°C, and a final extension step of 5 minutes at 72°C. Successful labeling of PCR2 products was then checked with the Fragment Analyzer Agilent 5300 using the DNF-910-33 dsDNA Reagent Kit (35–1,500 bp) protocol (Agilent Technologies, Santa Clara, CA, United States) and concentration was determined with PROSize 3.0 software. Using the QIAgility (Qiagen, Hilden, Germany), samples were pooled together at equimolar concentration. The pool was then cleaned using NucleoMag NGSBeads and the DNA concentration was quantified using Tapestation 4150 (Kit HSD 5000, Agilent Technologies, Santa Clara, CA, United States). The amplicon pool was sent to BaseClear (BaseClear B.V., Leiden, The Netherlands) for MiSeq Illumina sequencing (V3 2*300 PE platform).

### 16S rRNA gene metabarcoding data processing and analyses

The raw reads were initially processed by BaseClear B.V. for demultiplexing (using bcl2fastq version 2.20, Illumina), and filtering based on two quality controls (using Illumina Chastity filtering, and a PhiX control signal filtering). The resulting reads were then analyzed using the DADA2 workflow for the inference of Amplicon Sequence Variant (ASV) (Callahan et al., 2016a; 2017), using the “dada2” R package following the workflow described in Callahan et al., (2016b) and guidelines described in the online tutorial (https://benjjneb.github.io/dada2/tutorial.html). Filtering and trimming parameters were set as follows: truncation length of 270 bp and 240 bp for forward and reverse reads, respectively, maxN = 0, maxEE = 2, and truncQ = 2. After constructing the ASV table, chimeric sequences were filtered and removed and the taxonomic assignment was performed using the Silva v138 reference database (Quast et al., 2013). In alignment with the SILVA v138 taxonomy, the phylum Crenarchaeota was used instead of Thaumarchaeota which was reassigned at the class level as Nitrososphaeria (Parks et al., 2018). The ASV and taxonomy tables produced by the pipeline were then combined into a phyloseq object, together with the sample metadata table, using the “phyloseq” R package (McMurdie and Holmes, 2013). The dataset was subsequently filtered by removing all sequences from Eukaryota, chloroplast, and mitochondria (representing 0.2%, 0.1% and 0.9% of all reads, respectively). Data was then decontaminated with the negative controls and extraction blanks used as control samples, through the “decontam” R package (Davis et al., 2018). Rarefaction curves are plotted through the “vegan” R package (Oksanen et al., 2019). The *α*-diversity metrics were estimated using Chao1 (estimated richness), Pielou (evenness), and Shannon (both richness and evenness) indices on the rarefied dataset (rarefaction to the minimum library size, i.e. 10926 reads), using the “phyloseq” and “vegan” R packages (McMurdie and Holmes, 2013; Oksanen et al., 2019). According to the Shapiro tests, the diversity metrics significantly differed from the normal distribution. Consequently, the significance of the diversity metrics across the different groups of samples was investigated through non-parametric tests (Kruskal-Wallis followed by pairwise Wilcoxon tests) using the “agricolae” R package (de Mendiburu et al., 2019). Following recommendations for compositional approaches from (McMurdie and Holmes, 2014, Gloor et al., 2017), all other analyses were conducted without rarefaction, using the “phyloseq” R package, and the datasets normalized to the total number of sequences per sample (named “compositional dataset”). The *β*-diversity was analyzed with non-metric multidimensional scaling (NMDS) using Bray-Curtis dissimilarity. Differences in *β*-diversity between groups were statistically checked with one-way permutational multivariate analysis of variance (PERMANOVA) tests followed by pairwise Adonis tests, using the “vegan” R package. Differential analyses were performed using the “metacoder” R package (Foster et al., 2017), to identify the significant taxa differentially abundant according to the comparisons (i) between the two sponge classes, and (ii) among the photic zones for both classes separately. The differential analyses were performed with a subset of the compositional dataset excluding rare ASV (relative abundance < 0.04%).

### Extraction, sample preparation, and data acquisition for UHPLC-ESI-HRMS metabolomics

Samples of sponges tissues filled with 96% EtOH, were sonicated for 30 minutes. The extract was then concentrated with the CentriVap Benchtop Vacuum Concentrators (Labconco^TM^, Kansas City, MO, United States). Fresh 96% EtOH was added to the sponge tissues, and the sonication-evaporation procedure was repeated twice. Extracts were transferred to 1.5 ml glass vials and stored at -2 0°C until the sample preparation for metabolomics.

UHPLC-MS samples were prepared by solubilizing 2 mg of dried sample in 0.5 ml of methanol (HPLC grade, Sigma-Aldrich, St. Louis, MO, USA). Ten quality control (QC) samples were prepared by mixing all the samples at equimolar concentrations, in addition to ten extraction blanks, and two analytical blanks. Samples and blanks were injected in a random order, with a QC for every five injections. The samples were analyzed using a UHPLC-DAD-HRMS, UltiMate 3000 system (Thermo Scientific, Waltham, MA, USA) coupled to a QTOF-II mass spectrometer equipped with an electrospray ionization (ESI) source (Bruker, Bremen, Germany). The separation was performed using a 2.1 mm x 150 mm, 2.6 µm Kinetex C18 column (Phenomenex, Torrance, CA, USA) at 40 °C and a 0.3 ml/min flow rate. For samples, QC and MeOH, 1µL was injected and eluted with a gradient of 0.1% formic acid in water (A) and 0.1% formic acid in acetonitrile (B) starting at 5% B (0-30 min), 98% B (30-35 min) and equilibrated at 5% B for 5 min. The capillary voltage was set at 4000 V, the drying temperature at 350°C (0.3 ml/min); the nebulizer gas pressure at 2.0 bar, and the gas drying at 8.0 ml/min. The samples were analyzed in positive mode using a scan range of 100-1600 *m/z*.

### Metabolomics data processing, statistical analysis, and annotation

The UHPLC-MS/MS raw data files obtained were first converted into mzML files using the open-source msConvert tool from ProteoWizard library (http://proteowizard.sourceforge.net). The mass detection, chromatogram alignment, and ion deconvolution were performed using Metaboscape (Bruker Daltonics GmbH, Bremen, Germany, version 4.0) following the T-ReX 3D processing workflow (Version 1.9, detailed settings provided in Supplementary Information (SI, see material and method section). Briefly, the software performed automated mass calibration and deisotoping, followed by alignment of the resulting feature retention times using a LOESS-based alignment algorithm. The resulting feature table was filtered through an in-house R script (Paix et al., 2021), to remove successively experimental and analytical bias according to signal/noise ratio (using blanks), coefficient of variation (using QCs), and coefficient of correlation (using samples). The α-chemodiversity was analyzed through the data matrix normalized to the sum of the chromatographic peak areas, using the Shannon index and the vegan R package. Differences in α-chemodiversity within groups were tested through an ANOVA test, using the vegan R package. Multivariate analyses were analyzed using MetaboAnalyst 6.0 online web tool (Pang et al., 2024), through normalization to the sum of the chromatographic peak areas, a log10-transformation, and a Pareto scaling. Through a first Principal Component Analysis (PCA), QC samples were confirmed to be grouped, and blanks were identified as outliers. A second PCA was then conducted with the sponge samples only, to determine the overall clustering patterns of the holometabolome chemodiversity. Statistical differences in metabolome profiles between the sample groups were assessed with a PERMANOVA test and the Euclidean distance with the vegan R package. Following PCA, Partial Least Square Discriminant Analyses (PLS-DA) were conducted to reveal the most discriminant features involved in the differences between (i) the sponge class and (ii) the photic zones. Using retention time and MS/MS spectra, the annotation efforts were dedicated to these features with a VIP score > 2 for the class differences, and VIP score > 3 for the zone differences for both classes. These thresholds were defined to select only significant features, according to ANOVA and T-tests. The annotation of this discriminant feature list was facilitated by the Feature-Based Molecular Network (FBMN) to identify clusters of compounds sharing similar MS/MS spectra and fragmentation patterns. The mass spectrometry data were first processed with Metaboscape and the results were exported to Global Natural Products Social Molecular Networking (GNPS; Wang et al. 2016) for FBMN analysis. The data was filtered by removing all MS/MS fragment ions within ±17 Da of the precursor *m/z* MS/MS spectra were window filtered by choosing only the top 6 fragment ions in the +/- 50 Da window throughout the spectrum. Parameters used for building the FBMN within GNPS are detailed in the Supplementary Information (SI). The FBMN was finally analyzed using the Cytoscape software (version 3.10.2). The level of confidence in the annotation of the *m/z* features was determined according to Schymanski et al. (2014).

### Multi-omics integration of metabarcoding and metabolomics datasets

Shannon measures from both metabarcoding and metabolomics datasets were compared and their correlation was tested using the Spearman rank correlation using the “ggpubr” R package (Kassambara, 2023). A Procrustes analysis was performed using the metabolomics and metabarcoding dissimilarity matrices through the “vegan” R package. For the metabolomics, the dissimilarity matrix was calculated with the Euclidean distance and corresponded to the target matrix (X matrix). For the metabarcoding, the Bray-Curtis index was used to calculate the matrix to be rotated (Y matrix). Following the Procrustes analysis, a Mantel test was performed to confirm the significance of the correlation between the two dissimilarity matrices. Both datasets were also analyzed together into a supervised multi-omics analysis to assess the co-occurrences of the most discriminant ASVs and metabolites according to the discriminations between the depth zones and sponge classes. To conduct this approach, the DIABLO analysis was performed with the “mixOmics” R package through the *block.splsda*() function (Lê Cao et al., 2009; Rohart et al., 2017). The supervision factor gathered the samples according to their respective depth zone and sponge class (resulting in 5 groups: mesophotic demosponges, upper-rariphotic demosponges, lower-rariphotic demosponges, upper-rariphotic hexactinellid, and lower-rariphotic hexactinellid). An optimal number of six components was obtained based on the performance plot calculated for validation of the DIABLO model. The tuning step of the DIABLO pipeline resulted in optimal numbers of 25, 5, 5, 5, 5 ASVs and 6, 20, 30, 5, 30, 5 metabolites to retain for the first six components. Using this final DIABLO model, the resulting heatmap was obtained and plotted using the *cimDIABLO*() function, to visualize distinct clusters of samples gathered according to the co-occurrence of discriminant variables.

## Results

### Morphological and molecular identification of sponges and associated fauna specimens

The analysis of the demosponge samples identified 13 distinct species across 11 genera, eight families, and five orders (**Table S1, Figure S3**). The most diverse group was the order Tetractinellida with a total of nine species belonging to four families: *Aciculites higginsii* Schmidt, 1879 (family Scleritodermidae), *Gastrophanella implexa* Schmidt, 1879 (family Siphonidiidae), *Geodia* cf. *megastrella* Carter, 1876, *Geodia* aff. *curacaoensis* van Soest, Meesters, Becking, 2014, two unidentified *Geodia* sp., *Calthropella (Pachataxa) lithistina* (Schmidt, 1880), *Penares mastoideus* (Schmidt, 1880) (family Geodiidae), *Cinachyrella kuekenthali* (Uliczka, 1929) (family Tetillidae); followed by the order Haplosclerida; *Neopetrosia eurystomata* van Soest, Meesters, Becking, 2014; *Petrosia* aff. *weinbergi* and two unidentified *Petrosia* sp. (family Petrosiidae); order Biemnidae, *Biemna microacanthosigma* Mothes, Hajdu, Lerner & van Soest, 2004 (family Biemnidae); Order Scopalinida *Svenzea zeai* (Alvarez, van Soest & Rützler, 1998) (family Scopalinidae) and the order Suberitida: *Topsentia ophiraphidites* (de Laubenfels, 1934) (family Halichondriidae).

The analysis of hexactinellid samples revealed 7 distinct species across 7 genera, distributed among 5 families (**Table S1, Figure S3**). The most diverse family was Sceptrulophora *incertae sedis*, which included three species: *Lefroyella* sp., *Conorete pourtalesi* Reiswig & Dohrmann, 2014, and *Verrucocoeloidea liberatorii* Reiswig & Dohrmann, 2014. Additionally, each of the following families was represented by one species: *Heterotella* sp. nov (family Euplectellidae), *Myliusia* sp. (order Hexasterophora *incertae sedis)*, *Dactylocalyx pumiceus* Stutchbury, 1841 (family Dactylocalycidae), and *Hexactinella* sp (family Tretodictyidae).

The analysis of sponge-associated fauna revealed the presence of zoanthids colonizing three glass sponges: *C. pourtalesi*, *V. liberatorii,* and to a lesser extent *Hexactinella* sp. (**Figure S4**). For the three sponges, these zoanthids specimens corresponded to *Vitrumanthus schrieri* Montenegro & Reimer, 2022 based on their 28S rRNA gene sequences. Shrimp pairs were found inhabiting internal cavities of *C. pourtalesi* specimens, and were identified as *Eiconaxius caribbaeus* (Faxon, 1896) based on morphological analyses (pers. comm. Charles Fransen).

### Diversity and composition of prokaryotic communities

Rarefaction curves reached a plateau for all samples (**Figure S5A**), indicating a good coverage of the richness through the number of reads obtained for the whole study. Among the *α*-diversity metrics of the sponge-associated prokaryotic communities, the Pielou (evenness) and Shannon (richness and evenness) indices were significantly higher for Demospongiae compared to Hexactinellida samples (Kruskal-Wallis followed by post-hoc Wilcoxon tests: *p* < 0.001 for both indices, **Figure S5B**). For Chao1, no significant differences were observed when comparing the two sponge classes (post-hoc Wilcoxon test: *p* > 0.05, **Figure S5B**). For the three indices, no significant differences were observed between photic zones, either within Demospongiae or Hexactinellida (post-hoc Wilcoxon test: *p* > 0.05, **Figure S5B**).

The *β*-diversity, investigated through the Bray-Curtis dissimilarity index using an NMDS analysis, revealed two distinct class-specific clusters separated along the first axis with Demospongiae samples on the left side, and Hexactinellida samples on the right side (**Figures 1A** and **1B**). Differences between the two classes were confirmed to be significant through the two-way PERMANOVA tests followed by pairwise comparison tests (*p* < 0.01, **Tables S2** and **S3**). For the Demospongiae, the NMDS and PERMANOVA analyses also revealed distinct clusters between the three photic zones (**Figure 1A, Table S3**), while no differences were observed within the Hexactinellida samples. When depth is fitted as a continuous factor within the NMDS analysis, increasing gradients (from 100 to 300 m) can be distinguished (**Figure 1**) but towards opposite directions along the NMDS1 axis for each class (i.e. right side for Hexactinellida and left side for Demospongiae). Significant differences were observed when comparing the *β*-diversity among sponges genera (**Tables S4 and S5**). Clusters being specific to the sponge species clusters can be observed on the NMDS plot (**Figure 1B**), for example with *D. pumiceus* and *V. liberatorii* for the hexactinellids, as confirmed by the multivariate pairwise comparison test (**Table S5**). The NMDS plot displaying ASV scores (**Figure 1C**), revealed the presence of dominant archaeal ASVs (Crenarchaeota with relative abundances >1% in the overall dataset) associated with the Hexactinellida cluster. More precisely, the two most abundant ASVs; ASV1 and ASV2 (both with relative abundance >5%) were found to be specifically associated with the deepest rariphotic samples collected (∼250-300m), within the *V. liberatorii* and the *C. pourtalesi* cluster, respectively (**Figure 1B** and **Figure 1C**). Within the other Hexactinellida samples (*Mylusia*, *Dactylocalyx*, *Hexactinella*, *Lefroyella*), distinct ASVs from Crenarchaeota were found (e.g. ASV5, ASV6, ASV8), in addition to Proteobacteria (ASV3 and ASV4) and SAR324 (ASV7). These ASVs were found in lower abundance than ASV1 and ASV2 (ranging from 1% to 1.8%). ASVs found specifically associated with Demospongiae were observed in lower abundance (<1%), and mostly affiliated (i) to the PAUC34f, and Chloroflexi for the mesophotic and upper-rariphotic ones, and (ii) to Crenarchaeota for the lower-rariphotic ones.

**Figure 1.**
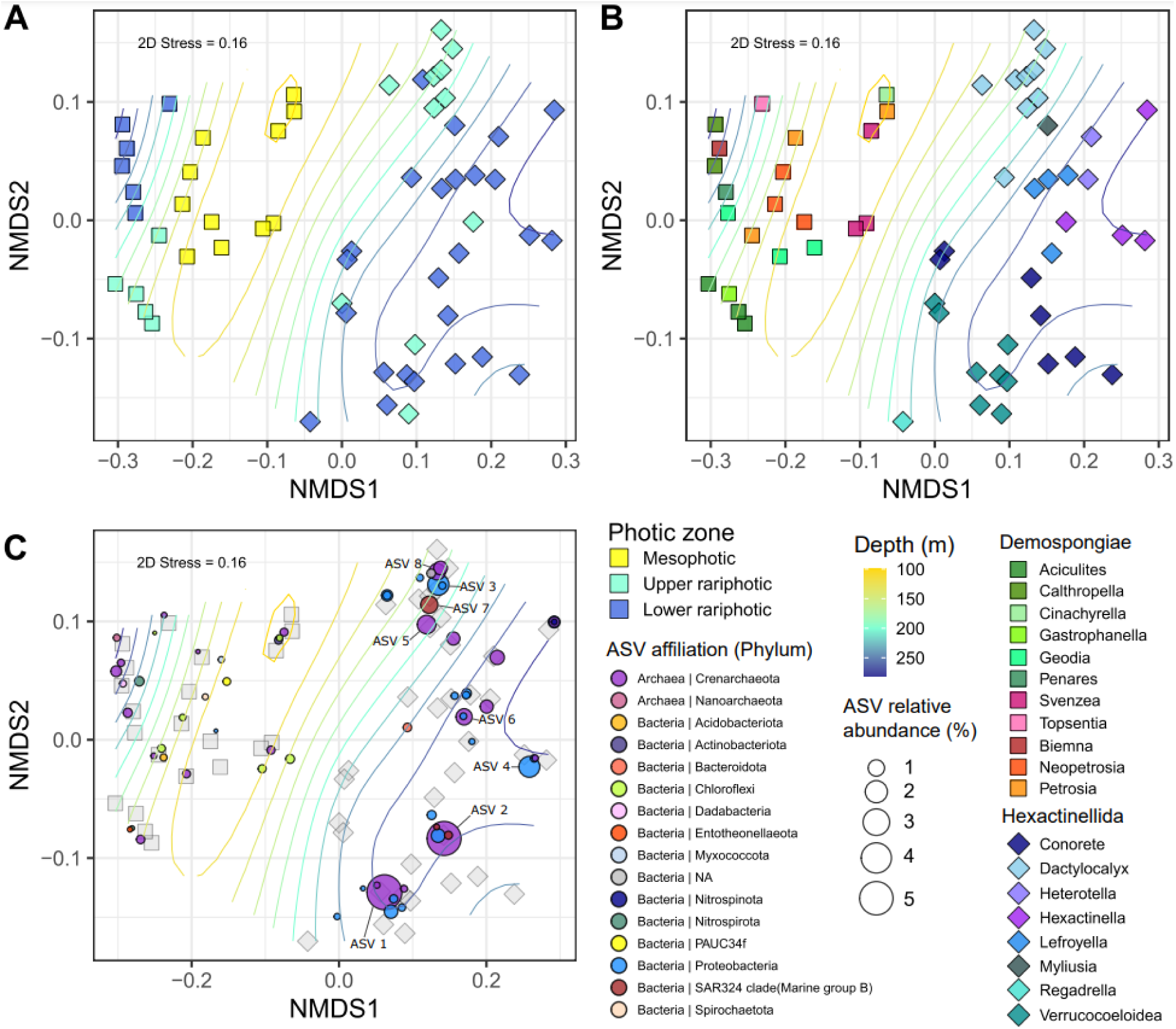
NMDS plots (built with the Bray-Curtis dissimilarity index) representing the *β*-diversity of the prokaryotic communities associated with the Demospongiae and Hexactinellida samples. **A** and **B**. Plots of the sample scores. **A**. Color codes were represented according to the photic zones. **B**. Color codes were represented according to the sponge genus. **C**. Plot of the ASVs scores. Color codes were represented according to the prokaryotic phyla. (A, B, and C) Curved lines represent the fitting of the depth gradient using the *ordisurf* function.

The prokaryotic community composition plotted at the family level confirmed the major differences observed between Demospongiae and Hexactinellida samples (**Figure S6**). Within the Demospongiae, Chloroflexi and Crenarchaeota were the two dominant phyla, represented by an unaffiliated family from the SAR202, and the Nitrosopumilaceae, respectively. To a lesser extent, other families were also dominant, such as Nitrospinaceae (Nitrospinota phylum), Nitropiraceae (Nitrospirota phylum), Nitrosococcaceae (Proteobacteria phylum), two unaffiliated families from the Acidobacteriota phylum, and an unaffiliated family from the PAUC34f phylum. For hexactinellids, higher intra- and inter-specific variation can be observed compared to demosponges (**Figure S6**). Among this heterogeneous composition within Hexactinellida species, Crenarchaeota and Proteobacteria were confirmed as dominant phyla. The Nitrosopumilaceae was the dominant archaeal family with a wide range of relative abundance varying from 1.24% to 83.75%, without clear host species or depth zone-specificity. Species-specific proteobacterial families can be distinguished, for example with an unaffiliated family of the UBA10353 group for *Hexactinella* sp., and the PS1 clade from the Parvibaculales for *V. liberatorii*. For archaea, host species-specificity could be discerned when analyzing their specific composition at the genus level (**Figure S7)**. More precisely, *Cenarchaeum* was the dominant genus of *C. pourtalesi* and *V. liberatorii,* while *D. pumiceus* and *Leyfroyella* sp. were found to harbor unaffiliated members of the Nitrosopumilaceae family. These latter were also dominant within the upper rariphotic Demospongiae, while the mesophotic sponges (except *C. kuekenthali* and one *N. eurystomata*) were dominated by Candidatus *Nitrosopumilus*.

The heat tree performed with all samples highlighted a large diversity of taxa differentially abundant between the two sponge classes across all taxonomic levels (**Figure S8**). The analysis confirmed our previous observations for the most abundant bacterial taxa found in significantly higher abundance within Demospongiae (**Figure S6**), with (i) Chloroflexi (Dehalococcoidia and Caldilineaceae), (ii) Nitrospirota (*Nitrospira* genus), (iii) Acidobacteriota (Vicinamibacterales), (iv) Entotheonellaeota (Entotheonellaceae). Within the Hexactinellida the bacterial taxa found significantly and differentially more abundant for this class were (i) the Planctomycetota (Phycisphaeraceae and Pirellulaceae), (ii) the Enterobacterales (*Vibrio* genus), (iii) the cyanobacterial genus Prochlorococcus MIT931, and (iv) the Bacteroidia (more specifically the *Fulvitalea* genus). Within the Crenarchaeotal family Nitrosopumilaceae, the Candidatus *Nitrosopumilus* genus was found to be significantly and differentially more abundant in Demospongiae, while *Cenarchaeum* was found specifically more abundant in Hexactinellida (**Figure S8**).

The differential pairwise analyses conducted with the Demospongiae samples revealed a large diversity of taxa being specific to each photic zone (**Figure 2**). Demospongiae sponges from the mesophotic zone were enriched with SAR202 and Caldilineaceae (both from the Chloroflexi phylum), Spirochaetota, and more specifically its genus *Spirochaeta*, Vicinamibacterales, and Candidatus *Nitrosopumilus.* Below the mesophotic, the families Microtrichaceae, Entotheonellaceae, Nitrosococcaceae, and Woeseiaceae were found to be more abundant within the upper rariphotic zone. In addition, the lower rariphotic demosponges were specifically enriched in Dadabacteriales, Thermoanaerobaculaceae, the TK10 cluster from the Chloroflexi phylum, and the *Cenarchaeum* from the Nitrosopumilaceae (**Figure 2**). For the Hexactinellida samples (**Figure S9**), a large diversity of taxa was also found differentially abundant for the upper and lower rariphotic zones respectively. More precisely, the upper-rariphotic Hexactinellida sponges were enriched in Leptospiraceae, *Prochlorococcus*, Rhodobacteraceae, SAR11 Clade Ia, *Coxiella*, while lower-rariphotic ones were enriched with *Vibrio*, *Fluvitalea*, Phycisphaeraceae and *Cenarchaeum* (**Figure S9**).

**Figure 2.**
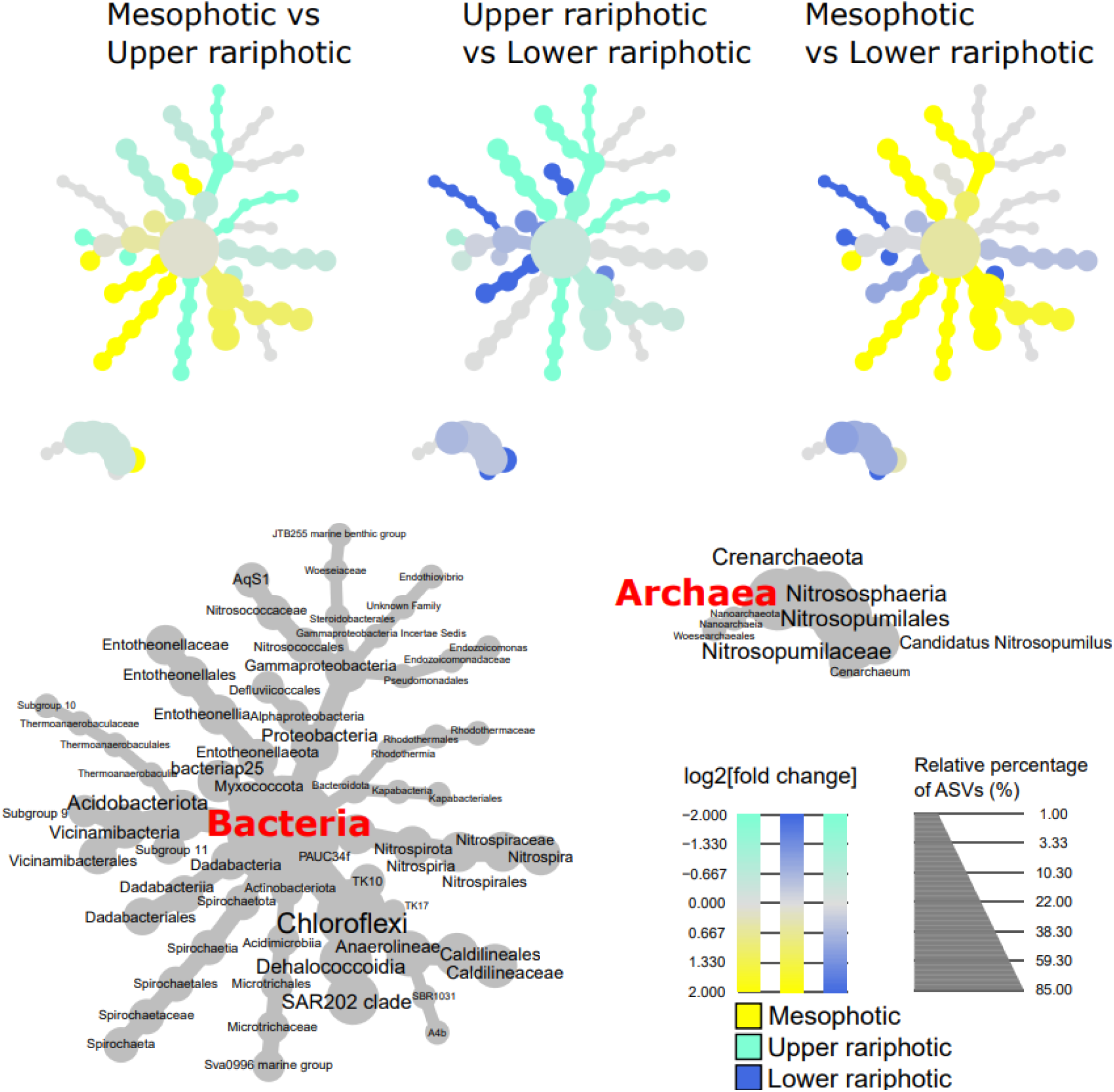
Phylogenetical heat trees performed with Demospongiae samples, and representing taxa significantly and differentially abundant between mesophotic, upper rariphotic, and lower rariphotic samples. For each taxon, (i) the colors of their associated nodes correspond to the log2 fold change between the ratio of the mean relative abundance within samples of each zone, (ii) and the size of their nodes corresponds to their relative abundance.

### Chemical diversity and annotation of sponge-holobiont metabolomes

Demosponge holometabolomes were characterized by a significantly higher *α*-chemodiversity (Shannon index) compared to their hexactinellid counterparts (ANOVA test: *p* < 0.05). However, no significant differences were observed among the photic zones (ANOVA test: *p* > 0.05, **Figure S5C**). PERMANOVA and pairwise comparison tests revealed significant differences between the metabolome profiles of the two sponge classes, as confirmed by the PCA plot (**Figures 3A**, **Tables S2 and S3**). For Demosponges, mesophotic and lower-rariphotic samples appeared clustered separately on the PCA plot. Significant differences were confirmed among the photic zones for this class, through the two-way PERMANOVA test followed by a pairwise comparison test (**Tables S2 and S3**). For hexactinellids, no clear differences among the photic zones or depth gradient can be observed, according to the PCA plots and tests (**Figure 3A, Table S2**). Within demosponges, the PCA plot did not reveal clear genus-specific clusters of holobiont metabolome profiles (**Figure 3B**). However, differences between species can be observed for hexactinellid samples, as also suggested by the PERMANOVA and pairwise comparison tests (**Figure 3B, Tables S4 and S5**).

**Figure 3.**
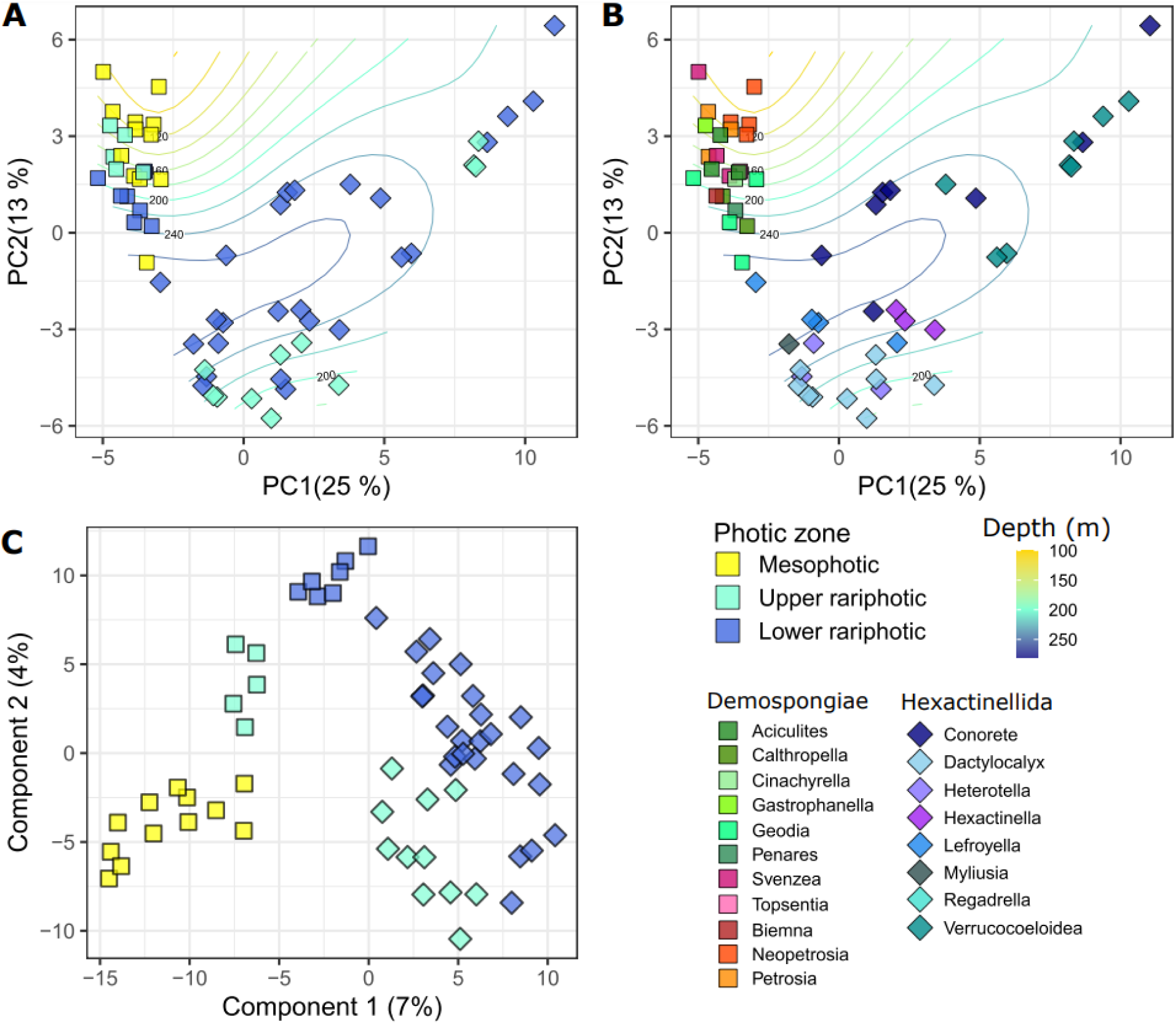
Plots of multivariate analyses of the metabolome associated with the Demospongiae and Hexactinellida holobionts. **A** and **B.** PCA plot of the metabolomes. Curved lines represent the fitting of the depth gradient using the *ordisurf* function. **A.** Color codes are defined according to the photic zones. **B.** Color codes are represented according to the sponge genus. **C**. PLS-DA plot conducted with the photic zone as a supervised factor.

PLS-DA were conducted to confirm the presence of metabolites significantly involved in the discrimination between the sponge class or photic zone, and target the annotation strategy specifically towards these compounds. In addition to the major differences between the sponge classes, discriminations within the photic zones were also observed with a continuous shift from the mesophotic to the lower-rariphotic zones (**Figure 3C**). PLS-DA performed with Demospongiae and Hexactinellida samples separately and with the photic zone as a supervision factor, also revealed discrimination between photic zones (**Figure S10**).

These PLS-DA allowed to establish a list of the most discriminant features (VIPs) between (i) Demospongiae and Hexactinellida (**Table S6,** VIP scores > 2), (ii) photic zones for the Demospongiae samples (**Table S7,** VIP scores > 3) and (iii) photic zones for the Hexactinellida samples (**Table S8,** VIP scores > 3). Annotation efforts were focused on these compounds, and facilitated through a first identification step of the main chemical families, using the molecular networking approach. The FBMN allowed the gathering of the MS/MS fragmented ions within distinct clusters associated with specific chemical families (**Figure S11**). The annotation was first focused on clusters gathering VIPs (**Figures S12 and S13)**. Cluster A represented the main cluster of the overall FBMN, gathering a total of 12 phosphatidylcholines (PC), together with 12 *lyso*-PCs corresponding to a specific sub-cluster (**Figure S12**). PC and *lyso*-PCs were annotated based on the characteristic fragment ion of phosphocholine at *m/z* 184.07 [C_5_H_15_NO_4_P]^+^ (Carriot, Paix et al., 2021). Fragment ions associated with the carbon chains allowed the identification of their acyl chain length and unsaturation number, for both PCs and *lyso*-PCs (**Figure S12**). Cluster G gathered compounds characterized by typical 1:2:1 isotopic patterns indicating the presence of two bromine atoms (**Figure S13**). The isotopic cluster at *m/z* 539.0407 and 541.0402 (corresponding to [M+H]^+^ and [M+H+2]^+^, respectively) matched with the chemical formula C_20_H_24_Br_2_N_6_O_2_. The MS/MS fragmentation patterns confirmed the identification of aphrocallistin, a dibrominated alkaloid previously isolated and characterized by Wright et al., 2009 from the glass sponge *Aphrocallistes beatrix* (**Figure S14, Table S6**). Other compounds from cluster G were linked to aphrocallistin with a very similar MS/MS fragmentation pattern (**Figure S13**). These compounds corresponded to aphrocallistin derivatives, such as *m/z* 569.0453 corresponding to the chemical formula C_21_H_26_Br_2_N_6_O_3_ (**Figure S12**). Cluster D was also found to share a similar mass spectrum to the dibrominated compounds of cluster G, especially with the identification of an aphrocallistin fragment ion, and a brominated purinone (*m/z* 286.0286, chemical formula: C_9_H_12_BrN_5_O). Cluster I gathered peaks with MS/MS spectra sharing a neutral loss of *m/z* 141.02 (**Figure S12**). This fragmentation pattern is characteristic of the (PEs), with the loss of the phosphoethanolamine moiety (Carriot, Paix et al., 2021; Godzien et al., 2015). A total of 2 PEs and 7 *lyso*-PEs were identified from this cluster. Finally, cluster O was characterized by features putatively annotated as amino acid derivatives (**Figure S11**, **Tables S7 and S8**).

Among the 28 first VIPs (VIP score > 2) discriminating demosponge from hexactinellid holometabolomes, 12 were putatively annotated as phospholipids, including 9 *lyso*-PCs (**Table S6**). Among them, *lyso*-PC(C17:0), *lyso*-PC(O-C17:0), *lyso*-PC(C17:1) and *lyso*-PC(O-C18:0) were found to have higher abundance within the Demospongiae samples (T-test: *p* < 0.001 for all, **Figure 4A**, **Table S6**). Conversely, *lyso*-PC(C20:4), *lyso*-PC(C18:0), *lyso*-PC(C20:5), *lyso*-PC(C19:0), and *lyso*-PC(C20:1) were found to discriminate the two classes with higher abundance within Hexactinellida (T-test: *p* < 0.001 for all, **Figure 4A**, **Table S6**). Additionally, the other phospholipids putatively identified as *lyso*-PI(C19:0) and *lyso*-PI(C18:0) were found as discriminant compounds with higher abundance in Demospongiae, while higher abundance within Hexactinellida was observed for *lyso*-PE(C20:4) (**Figure 4A, Table S6**). Aphrocallistin and one of its derivatives (VIP n°25 corresponding to C_21_H_26_Br_2_N_6_O_3_) were both characterized by a discriminant higher abundance in Hexactinellida sponges compared to Demospongiae (**Figure 4A**). These compounds also revealed distinct normalized concentrations within the different Hexactinellida genera, with higher concentrations observed within *V. liberatorii, C. pourtalesi,* and *Hexactinella* sp. (**Figures S15**). A compound identified as xanthurenic acid (**Figure S16**), was also discriminant between the two classes with significantly higher concentrations for Hexactinellida samples (**Figure 4A**), and more specifically within *C. pourtalesi* (**Figure S15**). Discriminant differences within photic zones for Demospongiae samples were observed for *lyso*-PC(O-C18:0), PC(O-C18:0/O-1:0), Gln-Lys and Gln-Pro with higher abundance within the lower rariphotic zone compared to the mesophotic one (**Figure 4B, Table S7**). Conversely, rhodosamine and salsolinol were found to be discriminant with relative concentration decreasing with the depth. Within Hexactinellida, the 17 first VIPs involved in the discrimination between the upper and lower rariphotic zones (VIP scores >2), were essentially identified as phospholipids and dipeptides (**Table S8**). VIPs annotated as *lyso*-PS(C15:1), *lyso*-PC(C22:5), *lyso*-PE(C22:2), *lyso*-PE(C22:4), *lyso*-PE(C20:4), Leu-Leu, and Leu-Val were discriminant with significantly higher abundance in the upper-rariphotic zone (**Figure 4C**, **Table S8**). Conversely, a higher abundance of xanthurenic acid was observed in the lower rariphotic zone, as *C. pourtalesi* were only found in this specific zone (**Figure 4C, Table S8**).

**Figure 4.**
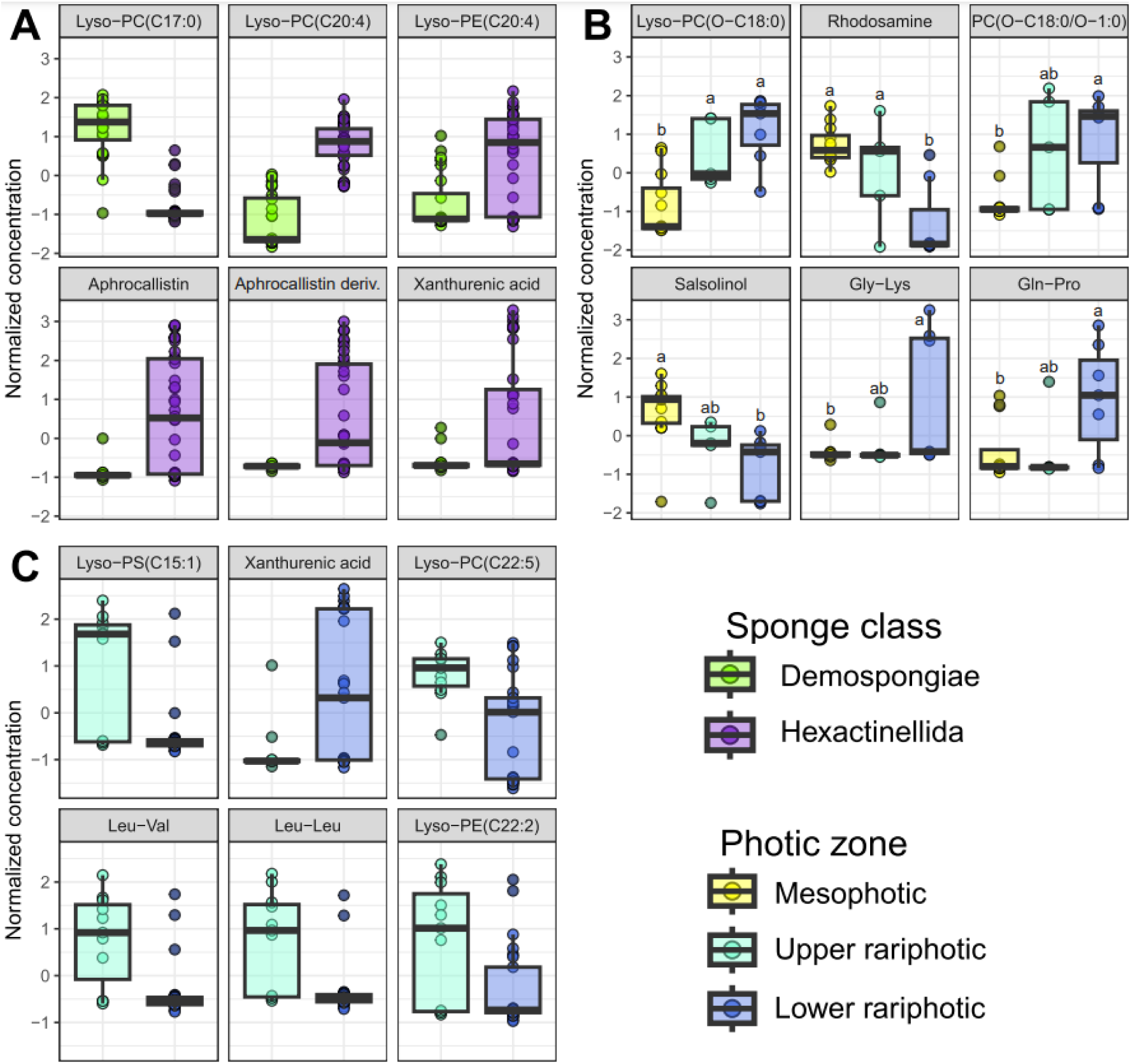
Normalized concentrations of discriminant metabolites putatively annotated with a confidence level ≥ 2. **A.** Discriminant metabolites (VIPs scores >2, *p* < 0.0001) according to the sponge classes. **B.** Discriminant metabolites (VIPs scores >3, *p* < 0.05) according to the photic zones for Demospongiae samples. Lowercase indices correspond to Tukey’s HSD pairwise comparison test. **C.** Discriminant metabolites (VIPs scores >3, *p* < 0.05) according to the photic zones for Hexactinellida samples. Results from the ANOVA and T-tests are presented in Tables S6, S7 and S8.

### Multi-omics analyses of metabarcoding and metabolomics datasets

A weak but significant linear correlation was observed between the Shannon measures of both datasets (Pearson correlation: r = 0.34, *p* = 0.011). This correlation was mainly driven by the differences between the two sponge classes (**Figure 5A**). Similarly, a significant correlation was also confirmed between the distance matrices of both datasets (Mantel tests: r = 0.22, *p* = 0.002), being also driven by sponge class differences (**Figure 5A and 5B**). Additionally, *V. liberatorii* and *C. pourtalesi* samples appeared to be clustered together, and separately from the rest of the other hexactinellids samples (**Figure 5C**). The heatmap built through a multi-omics supervised analysis (DIABLO), revealed a total of 135 co-occurring discriminant features (44 metabolites and 91 ASVs) involved in the differences between the samples grouped in their respective class and photic zones. The hierarchical clustering confirmed the major differences between the two classes, but also the differentiation between the photic zones. Chloroflexi and Actinobacteriota ASVs were clustered together with ether-linked *lyso*-PCs and derivatives of amino acids and amino sugars (e.g. dipeptides and rhodosamine), with higher abundance in the demosponges holobionts. A second cluster sharing ASVs and metabolites being differentially more abundant in the *C. pourtalesi* samples revealed *Cenarchaeum* and SAR324 Marine group B ASVs, together with primary lipids and the xanthurenic acid. Finally, hexactinellids were characterized by a higher abundance of several long-chain *lyso*-PCs [e.g. *lyso*-PC(C20:4) and *lyso*-PC(C20:5)], and aphrocallistins as previously observed in the VIP analyses.

**Figure 5.**
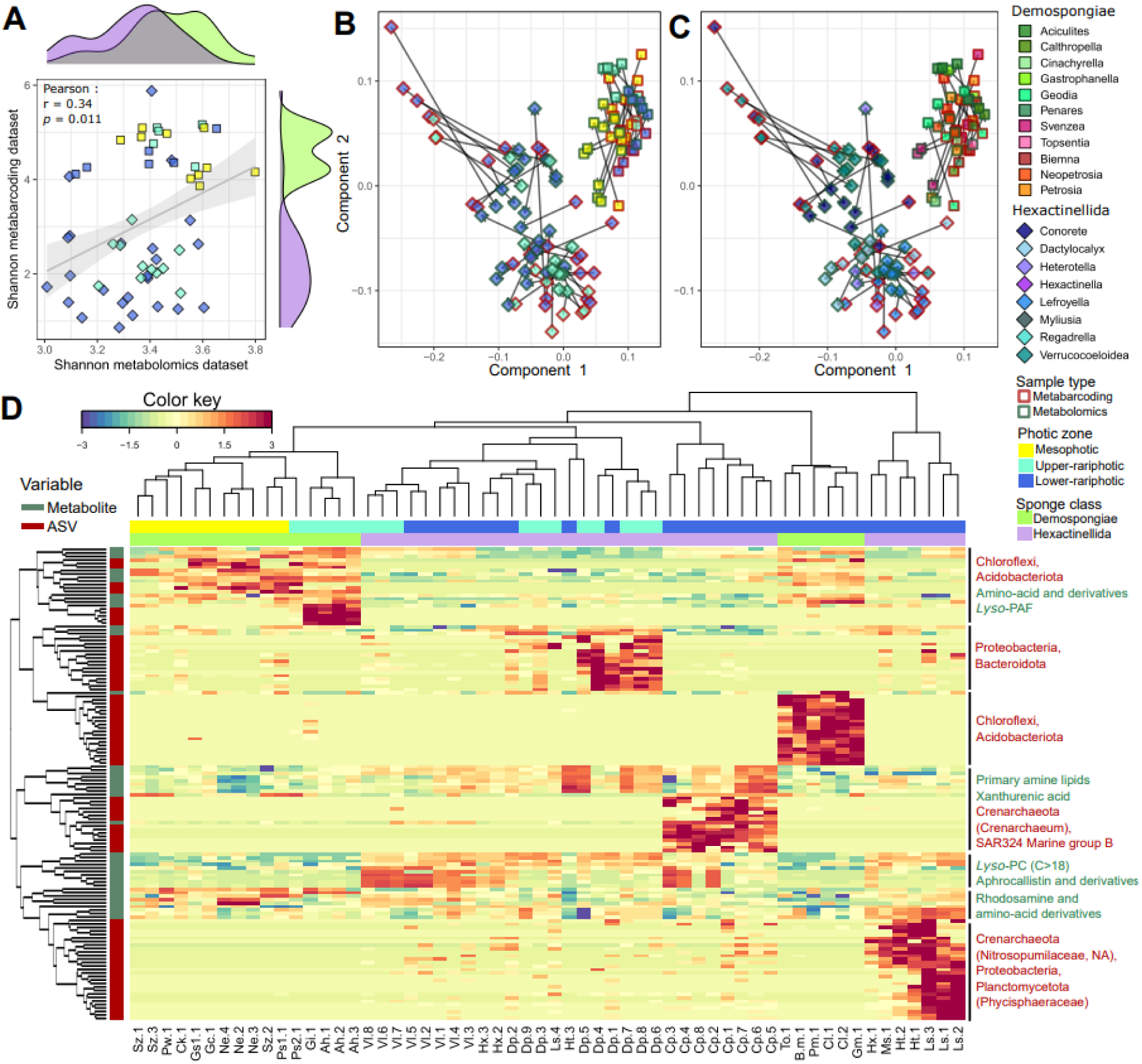
Multi-omics analysis of the sponge holobionts integrating metabarcoding and metabolomics datasets. **A.** Comparison of Shannon indices obtained through the metabarcoding and metabolomics dataset. **B.** Plot of the Procrustes analysis performed with both datasets, displaying the color codes for the photic zones. **C.** Plot of the Procrustes analysis performed with both datasets, displaying the color codes for sponge genera. **D.** Heatmap of the supervised multi-omics analysis (DIABLO). Sample names and abbreviations are listed in Table S1.

## Discussion

### Major differences in host-microbiome adaptive strategies are observed between the sponge classes

Whether analyzed independently or through a multi-omics analysis, the overall holometabolome profiles and prokaryotic community structure revealed major differences between sponge classes. Similar patterns have been observed in deep-sea sponge microbiomes in the South Pacific Ocean off the coast of New Zealand (Steinert et al., 2020), and more broadly in the Atlantic Ocean (Busch et al., 2022). In our study, all demosponge samples exhibited a High Microbial Abundance (HMA)-like community composition dominated by the phyla Chloroflexi, Acidobacteriota, and PAUC34f (Moitinho-Silva et al., 2017). In contrast, hexactinellid samples displayed lower evenness and a LMA-like structure, primarily dominated by a few Gammaproteobacteria and Crenarchaeota ASVs. These class-specific differences in microbial composition and abundance, emphasize the need to consider distinct host-microbiome adaptive strategies in deeper marine environments. Notably, hexactinellids also exhibited lower chemical *α*-diversity (Shannon index), which might be explained by a lower microbial richness and consequently a lower diversity of microbial-derived metabolites.

The relationships between the datasets, based on *α*-diversity measures (Shannon index measures) and multivariate analyses (Mantel test, Procrustes, and DIABLO analyses) appeared primarily driven by class-level differences. This finding suggests limited functional redundancy between demosponge and hexactinellid microbiomes, but indicates partial redundancy within each class. Using a multi-omics approach that combined metabarcoding and metabolomics, similar patterns were observed at lower taxonomic levels in four Mediterranean sponge holobionts, where intraspecific correlations were weaker than interspecific ones (Mazzella et al., 2024). Functional redundancy has also been proposed in metagenomics studies of HMA sponges from Caribbean coral reefs (Lesser et al., 2022), and in metatranscriptomic analyses of deep-sea sponges (Diez-Vives & Riesgo, 2024). In this latter study, functional redundancy was suggested to contribute to the holobiont stability, supporting adaptation to extreme environments. However, the functional redundancy concept in marine microbiomes is increasingly debated, as many microbial functions remain uncharacterized or underestimated (Galand et al., 2018). Therefore, evaluating the correlation between overall taxonomical and functional diversity as done in our study, offers a crucial “annotation-free” approach to investigate functional redundancies beyond reliance on a limited set of known functions (Galand et al., 2018).

Clear differences in the overall variance of microbiome *β*-diversity and holometabolome patterns were observed across depth zones for demosponges. In contrast, for hexactinellids, which are restricted to depths below the mesophotic zone **(Figure 6A**), no distinct separation between the upper and lower rariphotic zones was evident. However, supervised analyses revealed a wide diversity of discriminant metabolites and prokaryotic taxa specifically associated with these depth zones in both sponge classes. We propose that these microbial and chemical markers offer valuable insights into the adaptive strategies of sponge holobionts in response to depth-related environmental factors such as temperature, light, pressure, nutrients, oxygen, and biotic interactions **(Figure 6B** to **6H**). A critical aspect of this study focused on the MS/MS annotation of these discriminant metabolites, to uncover their potential biosynthetic origins and ecological roles using literature reviews and reference databases (e.g. Lotus, MarineLit). While untargeted metabolomics can provide valuable information to supplement phylogenetic studies of glass sponges (Dohrmann et al., 2023), this research presents the first untargeted metabolomics dataset for hexactinellid samples investigated through annotation efforts.

**Figure 6.**
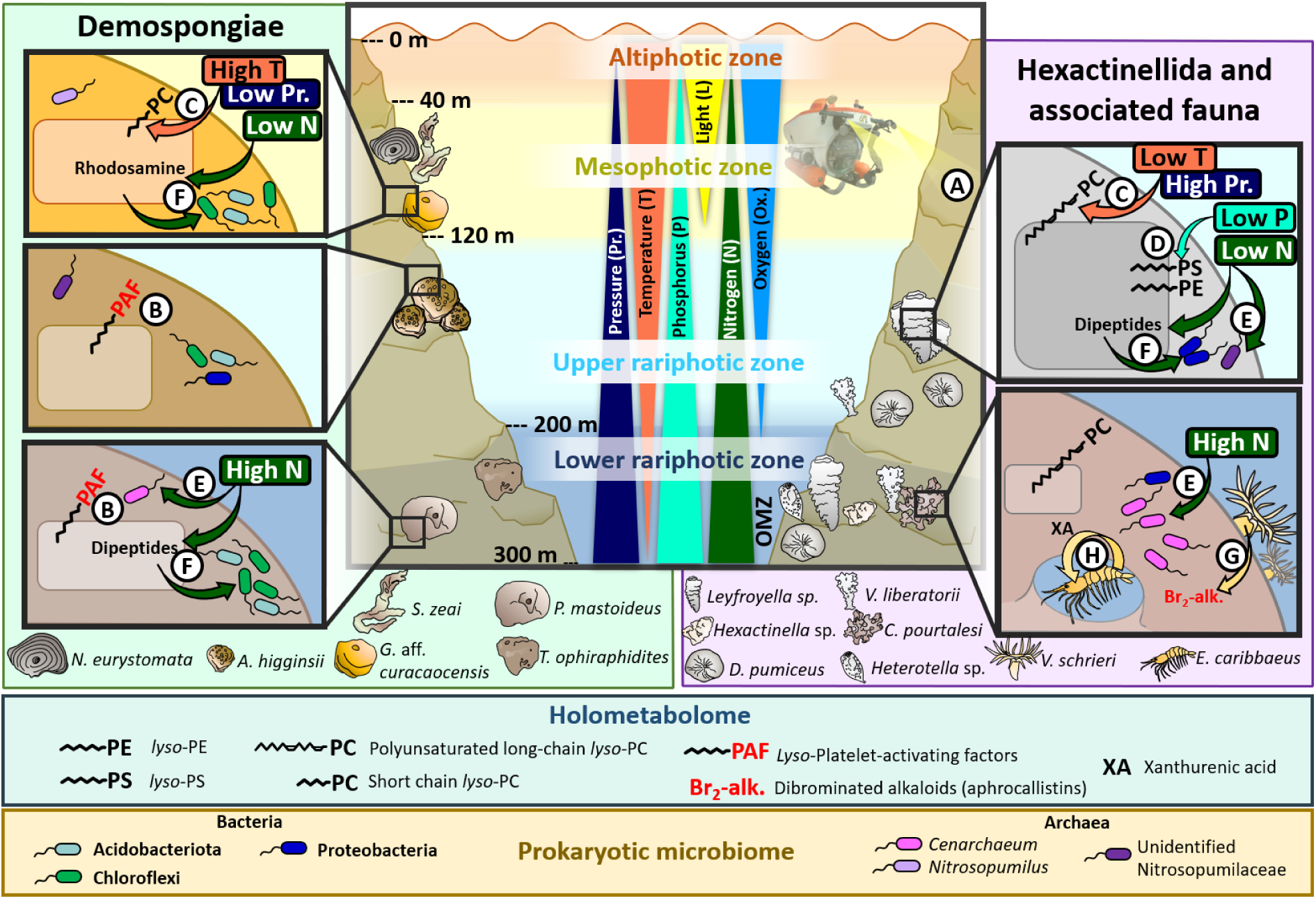
Summary of the distinct adaptive strategies of sponge holobionts collected within the mesophotic and rariphotic zones. **A.** No hexactinellids were occurring in the mesophotic zones, suggesting a preferential adaptation up to the lower rariphotic zone. **B**. *Lyso*-PAF are produced as immunolipids in the rariphotic demosponges. **C**. Homeoviscous adaptation of the *lyso*-PCs: the chain is longer under lower temperatures and higher pressure within hexactinellids, to increase the membrane fluidity. The opposite situation is observed in most of the demosponges holophospholipidome rather adapted to shallower conditions. **D**. The diversity of phospholipids synthesis pathways is suggested to increase as an adaptation to overcome the lower P-concentrations, leading to more diverse PLs classes. **E**. Within hexactinellids, the AOA members of the Nitrosupumilacae family are host-specific at the sponge species level and could be vertically transmitted. AOA symbionts such as *Cenarchaeum* are expected to ensure ammonia detoxification through its oxidation, under the OMZ. **F**. Amino sugars (e.g. rhodosamine) and amino acid derivatives (e.g. dipeptides) constitute a major source of DON for microbial members of the sponge holobionts, including the Chloroflexi. **G**. The zoanthid *Vitrumanthus schrieri* produces dibrominated alkaloids from the aphrocallistin family acting as chemical defense for the sponge hosts *C. pourtales*i, *V. liberatorii* and *Hexactinella* sp. **H**. The shrimp *Eiconaxius caribbaeus* inhabiting cavities of the hexactinellid *C. pourtalesi* might produce xanthurenic acid as an endogenous molting inhibitor to keep their body size small enough, maintaining their adult life cycle within the sponge. Metabolites in red correspond to putative chemical defenses (*lyso*-PAF as immunolipids, and aphrocallistin as brominated alkaloids).

### Temperature, pressure, and phosphate availability shape distinct phospholipidome adaptations

Phospholipids (PLs) and *lyso*-PLs (LPLs) emerged as the most diverse group of compounds identified in the sponge holometabolome extracts, including various forms of PCs, *lyso*-PC, PE, *lyso*-PE, and *lyso*-PS. These phospholipid classes are widely reported as dominant membrane components in many sponges, though their relative abundance can vary greatly between species (Genin et al., 2008; Mazzella et al., 2024). Among these membrane lipids, PLs and LPLs are also essential roles in sponge adaptation to environmental changes (Genin et al., 2008). Specifically, LPLs function as membranes-signaling metabolites, involved in various cell processes including growth, proliferation, survival, and apoptosis (Torkhovskaya et al., 2007). In sponges, LPLs may contribute to embryogenesis and morphogenesis (Ivanišević et al., 2011, Reverter et al., 2018). Among the annotated PLs and LPLs of our study, several were characterized as ether-linked *lyso*-PCs, PCs, and PEs, such as *lyso*-PC(O-C17:0), *lyso*-PC(O-C18:0) and PC(O-C18:1/O-C1:0). These lipids, previously identified in other marine sponges (Reverter et al., 2018, Ivanisevic et al., 2011, Zhang et al., 2022), are also named “platelet-activating factors” (PAFs). PAFs are known for their wide range of biological activities in marine invertebrates (Sugiura et al., 1992; Mannochio-Russo et al., 2023, Müller et al., 2004). In shallow water sponges, PAF production has been linked to immune responses, functioning as immunolipids in the antibacterial defense mechanism of *Suberites domuncula* (Müller et al., 2004). Additionally, PAFs may serve also as antifouling agents deterring settlement by ascidians, bryozoans, barnacles, algae, and mussels as observed in the Australian sponge *Crella incrustans* and the Mediterranean sponge *Oscarella tuberculata* (Butler et al., 1996, Ivanisevic et al., 2011). In this study, *lyso*-PC(O-C17:0) and *lyso*-PC(O-C18:0) were specifically associated with Demospongiae samples, and higher concentrations of *lyso*-PC(O-C18:0) and PC(O-18:1/O-1:0) detected in rariphotic sponges (**Figure 6B)**. While the ecological role of PAFs in the rariphotic sponges remains unclear, further research is necessary to explore their potential function as immunolipids, particularly in shaping host-specific and depth-specific microbial communities.

Interestingly, *lyso*-PCs identified as biomarkers for demosponges contained either one or no unsaturation and short acyl chains (below C17). In contrast, the other *lyso*-PCs identified as hexactinellid biomarkers exhibited a higher unsaturation degree (up to five double bonds) and longer acyl chains (from C18 to C20). We propose that the membranes of hexactinellid sponge species, or their associated bacteria, may be specifically adapted to the rariphotic zone, by producing long-chain polyunsaturated *lyso*-PCs. This adaptation likely mitigates the negative effects of lower temperatures and higher pressures on the membrane fluidity (**Figure 6C**). This process, known as homeoviscous adaptation, has been widely observed across diverse deep-sea phyla, including Chordata, Arthropoda, Cnidaria and Ctenophora (Cossins and Macdonald, 1989; Parzanini et al., 2018; Winikoff et al., 2024), as well in the shallow water phototrophic sponge species *Phyllospongia foliascens* (as *Carteriospongia foliascens* in Bennett et al., 2018).

Nutrient concentrations, including phosphate, are known to increase with depth reaching a maximum between 500-1000m. This pattern results from higher nutrient demand in the oligotrophic euphotic zone, and the downward transport of particulate organic matter into the deep ocean *via* the biological pump (Paytan & McLaughlin, 2007). As a result, varying phosphorus availability may drive distinct phosphorus cycling strategies in sponge-holobionts leading to differences in phospholipid composition, a pattern already observed in planktonic communities (Popendorf et al., 2011). In this study, hexactinellids in the upper-rariphotic zone exhibited a more diverse composition of phospholipid classes, with elevated concentrations of *lyso*-PE and *lyso*-PS in addition to the *lyso*-PC. This finding suggests that sponge-holobionts might diversify their phosphorus-uptake pathways under conditions of low phosphorus availability by producing multiple PL classes (**Figure 6D**).

### Ammonia oxidation and DOM cycling as key pathways of the N-metabolism within mesophotic and rariphotic depths

The nitrogen metabolism homeostasis of sponge holobionts is regulated by diverse prokaryotic symbionts involved in key nitrogen cycling processes, including nitrogen fixation, nitrification, denitrification, and anaerobic ammonium oxidation processes (Zhang et al., 2019). Among these, Ammonia Oxidizing Archaea (AOA) and Nitrite Oxidizing Bacteria (NOB) are particularly well-adapted and dominant within cold-water and deep-sea sponge holobionts (Tian et al., 2016; Wang et al., 2022; Steinert et al., 2020), playing crucial roles in the nitrification process (Glasl et al., 2024). However, there is no clear consensus regarding the species-specificity of these symbionts in deep-sea sponges. For instance, weak species-specificities were observed among samples from the South Pacific Ocean and Cantabrian Sea, suggesting a random acquisition of archaeal symbionts (Steinert et al., 2020) and the prevalence of functional redundancy within the microbiome (Diez-Vives and Riesgo, 2024). Conversely, Amplicon Sequence Variants (ASVs) from the AOA Nitrosupumilaceae family were identified as species-specific in deep-sea demosponges and hexactinellids from the Campos Basin, Southern Brazil) implying potential vertical transmission of these symbionts (Garritano et al., 2023). In shallow water sponges, vertical transmission has also been suggested. For example, nitrifying symbionts such as Candidatus *Nitrosokoinonia* and Candidatus *Nitrosymbion* have been identified in *Coscinoderma matthewsi* (Glasl et al., 2024), and the AOA *Cenarchaeum symbiosum* has been found across various sponge species in the Western Pacific, Caribbean, and Mediterranean Sea (Steger et al., 2008; Polónia et al, 2015, 2021). Consistent with the findings of Steinert et al., (2020), our study confirmed significant heterogeneity in the relative abundance of Crenarchaeota, within hexactinellid species. However, this observation is based on compositional data at the phylum and family levels, where relative abundances were analyzed with the bacterial community. When examining archaeal composition at the genus and ASVs level, species-specific patterns became evident. For example, *Cenarchaeum* was represented by two distinct ASVs, each predominantly associated with *C. pourtalesi* and *V*. *liberatorii*, respectively. Additionally, other ASVs from an unidentified *Nitrosopumilaceae* genus were specifically occurring within *D. pumiceus* and *Leyfroyella* sp. These findings suggest that the hexactinellid-AOA association is likely driven by a deterministic process, such as a vertical acquisition of a dominant host-specific nitrifying symbiont. However, once these symbionts are established their proliferation and relative abundance may be heavily influenced by external stochastic factors resulting in variability within the overall prokaryotic community.

Similar to phosphorus concentrations, variations in nitrogen availability from the mesophotic to the lower rariphotic depths may also be a critical factor influencing the nitrogen metabolism of both demosponges and hexactinellids holobionts. Within the microbiomes of both classes, the AOA genus *Cenarchaeum* was specifically associated with the lower rariphotic, suggesting a preference for habitats with elevated ammonium concentrations. This genus is represented by the species *Cenarchaeum symbiosum,* the first archaeal symbiont identified in sponges, originally discovered in the marine sponge *Dragmacidon mexicanum* (as *Axinella mexicana* in Preston et al., 1996). This widespread sponge symbiont is known for its role in ammonia detoxification through ammonia oxidation (Turque et al., 2010; Hallam et al., 2006; Bayer et al., 2020). We propose that *Cenarchaeum* plays a crucial role in supporting lower-rariphotic sponge holobionts by facilitating ammonia detoxification, particularly under the challenging conditions of the oxygen minimum zone (OMZ), which occurs at similar depths (**Figure 6E**).

Our metabolomics analysis revealed a diverse range of amino acids and their derivatives (e.g. dipeptides) and amino sugar (e.g. rhodosamine) that varied in normalized concentrations across photic zones in both demosponges and hexactinellids. These compounds are critical components of dissolved organic nitrogen (DON) in coral- and algae-dominated reef ecosystems (Fiore et al., 2015, Thobor et al., 2024). Amino acid transporters have been identified in several bacterial metagenomes of sponge symbionts, enabling the uptake of a wide diversity of carbon and nitrogen sources potentially produced by the sponge host (Moitiho-Silva et al., 2017; Botté et al., 2019; Podell et al., 2020; Taylor et al., 2021). Additionally, amino acids can be synthesized by the sponge host and subsequently metabolized by symbionts, as suggested by a transcriptomics analysis on *Xestospongia muta* (Fiore et al., 2015). Our multi-omics analysis highlights a co-occurrence of dipeptides and amino sugars together with several Chloroflexi and Acidobacteriota ASVs in demosponges. Notably, Chloroflexi (including the SAR202) are frequently implicated in the dissolved organic matter DOM cycling across shallow (Bayer et al., 2018, Maggioni et al., 2023), subtidal caves (Cleary et al., 2024b), mesophotic (Cleary et al., 2024a) and deep-sea ecosystems (Busch et al., 2020; Campana et al., 2021; Landry et al., 2017). These co-occurrences suggest that these microbial groups may play a key role in the cycling of amino acids and amino sugars, contributing to the DON pool within demosponges holobionts across all depth zones (**Figure 6F**). Within the dipeptide family, higher concentrations were detected in lower rariphotic demosponges and upper-rariphotic hexactinellids (**Figure 6F**).

Further research is needed to explore how these dipeptide distribution patterns, both class- and zone-specific, may reflect distinct strategies for DON cycling in response to the seawater chemistry unique to each zone.

### Symbiotic strategies between rariphotic hexactinellid-holobionts and their associated fauna revealed by in-depth annotation of secondary metabolites

In addition to phospholipids and dipeptides, another major compound family identified in this study was dibrominated alkaloids, unexpectedly detected as biomarkers for hexactinellids. Typically, Demospongiae and their microbiome are well known for producing diverse halogenated compounds (e.g. Turon et al., 2000; Kunze et al., 2013; Lira et al., 2011; Ueberlein et al., 2017; Agarwal et al., 2017; Bayona et al., 2021), but the biosynthesis of such compounds in Hexactinellida sponges has not been reported until now. Based on fragmentation pattern elucidation, this family of dibrominated alkaloids was identified as aphrocallistin and its derivatives. Aphrocallistin was initially isolated from extracts of the glass sponge *Aphrocallistes beatrix* (Wright et al., 2009), but subsequent studies suggested it was produced by zoanthids colonizing the sponge specimens (Young et al., 2021). Our findings support this hypothesis, as aphrocallistin and its derivatives were detected in lower rariphotic Hexactinellida species that were specifically colonized by zoanthids. These species were *V. liberatorii* and *C. pourtalesi*, and to a lesser extent *Hexactinella* sp. (**Figure 6G**). These zoanthids were identified as *Vitrumanthus schrieri* across all three glass-sponge species. The identification aligns with previous findings of *V. schieri* associated with hexactinellids (*V. liberatorii* and *Cyrtaulon sigsbeei*) collected in Curaçao (Reiswig & Dohrmann, 2014; Kise et al., 2022). Interestingly, Caribbean specimens of *Aphrocallistes beatrix* have also been reported in association with the zoanthid *Vitrumanthus vanderlandi* (Kise et al., 2022). This suggests that the production of aphrocallistin may not be limited to a specific zoanthid species, but could be a characteristic of the genus *Vitrumanthus*. Recent studies highlight that the extensive diversity of deep-sea zoanthids living as epibionts of Hexactinellida sponges remains largely unexplored (Hajdu et al., 2017; Kahn et al., 2020; Kise et al., 2022). These zoanthid-hexactinellid associations are often considered a form of symbiosis (Kise et al., 2022), and we propose that the production of aphrocallistins by zoanthids may serve as a source of chemical defense for the sponge host. In contrast, demosponges are well-known for hosting diverse bacterial symbionts with halogenation and dehalogenation activities (Ollinger et al., 2021), differentiating their chemical defense strategies from those of hexactinellids. For example, Cyanobacteria and Chloroflexi symbionts are known to participate in the bromination of inactive alkaloid precursors produced by the sponge host, leading to unique biosynthetic pathways for chemical defenses at the holobiont level (Turon et al., 2000, Sacristán-Soriano et al., 2011). However, in this study, the absence of Chloroflexi in Hexactinellida, which are instead dominated by Ammonia-oxidizing Archaea (AOA), and the low abundance of Cyanobacteria, particularly in the lower rariphotic zone, suggests that such bromination pathways are likely absent in hexactinellid holobionts such as *C. pourtalesi*. As a result, the symbiotic association with zoanthids producing bioactive dibrominated alkaloids may serve as an alternate chemical defense strategy for the hexactinellid sponges in deeper ecosystems (**Figure 6G**). In exchange, these heavily encrusting zoanthids (Kise et al., 2022) may benefit from the natural filter-feeding activity of the sponge to support their own nutrition. The functional role of the microbiome in *C. pourtalesi* and *V. liberatorii* both dominated by distinct *Cenarchaeum* symbionts remains to be fully explored. However, the production of aphrocallistin by zoanthids appears to be independent of the sponge’s microbial community structure. For instance, *A. beatrix* was dominated by the AOA Candidatus *Nitrosoabyssus spongiisocia* and the gammaproteobacterial Candidatus *Zeuxoniibacter abyssi* (Garritano et al., 2024), and yet still associated with these zoanthids.

In addition to aphrocallistins, the detection of xanthurenic acid (XA) provides another example of secondary metabolites involved in hexactinellid-fauna associations (**Figure 6H**). Specifically, XA has been previously isolated from extracts of the Antarctica sea sponge *Isodictya erinaceae* and identified as an endogenous molt inhibitor in crustaceans, such as the amphipod *Pseudorchomene plebs* (Vankayala et al., 2011), as well as in various crab species (Ohnishi and Naya, 1994; Naya et al., 1989a,b). Our results revealed higher concentrations of XA in *C. pourtalesi* samples. Notably, several specimens of this sponge species were found to harbor shrimp pairs within their internal cavities during subsampling. Based on morphological analysis, these shrimp were identified as *Eiconaxius caribbaeus* (Faxon, 1896 pers. observ. Charles Fransen). Adult shrimp of the genus *Eiconaxius* are known to exclusively inhabit the internal canals and cavities of deep-sea hexactinellid sponges (Kou et al., 2020). Therefore, it is plausible that these shrimps produce XA as an endogenous hormone to regulate their body size, enabling them to remain small enough to occupy the sponge’s internal cavities, which are approximately 5 mm in diameter.

Results from our untargeted metabolomics approach highlight the importance of comprehensive annotation strategies when analyzing complex sponge holometabolomes, as well as those of other marine models (Reverter et al., 2020; Carriot, Paix et al., 2021). These efforts have revealed diverse symbiotic strategies uncovering potential interactions between sponge holobionts and their associated fauna. These findings exemplify the "nested ecosystems" concept, emphasizing the need to integrate multiple ecological scales to fully understand the intricate relationships within sponge-holobionts and their interactions with the broader environment (Pita et al., 2018). By identifying a unique diversity of secondary metabolites within hexactinellid holobionts and their associated fauna, our study underscores the need for deeper exploration of chemical interactions within rariphotic assemblages. Such investigations pave the way for the discovery of new natural products, as demonstrated by the potential of aphrocallistin in cancer treatment research (Wright et al., 2009; Wright et al., 2011). Moreover, these findings highlight the critical importance of preserving these unique ecosystems, through dedicated conservation efforts (Stefanoudis et al., 2019).

## Conclusion

Our study revealed distinct adaptive strategies of sponge holobionts across and below the mesophotic zone, driven by host-microbiome interactions, depth-related environmental factors, and the production of unique secondary metabolites.

Using a multi-omics approach, we identified significant differences in microbial community structure and holometabolome diversity between Demospongiae and Hexactinellida sponges, indicating limited functional redundancy between these classes. Consistent with findings in other benthic communities, our results also support a clear delineation between mesophotic and rariphotic zones for demosponges, reflecting distinct adaptive strategies at the holobiont level. Depth-specific patterns in microbial and chemical markers (such as membrane phospholipid adaptations and nitrogen metabolism processes) underscore the influence of environmental gradients (e.g. temperature, pressure, and nutrient availability) in shaping these holobionts. Furthermore, the detection of secondary metabolites such as aphrocallistin and xanthurenic acid, which are associated with symbiotic fauna, reveals novel ecological strategies in rariphotic hexactinellids. These findings highlight the intricate symbiotic relationships between sponge holobionts and their associated fauna, supporting the concept of sponge holobionts as integral components of a nested ecosystem.

## Declarations

### Data availability

16S rRNA gene sequences for metabarcoding were deposited and are publicly available in the NCBI Sequences Read Archive (SRA) under the BioProject ID PRJNA1216067, accession number. MS data are available in the Mass Spectrometry Interactive Virtual Environment (MassIVE) repository MSV000096962 (reviewer password: ECG123I54HcdyoaT). FBMN workflows can be accessed in GNPS with the following links: https://gnps.ucsd.edu/ProteoSAFe/status.jsp?task=b9d95e6d74bd4428abe42b5667028897.

The R scripts used for all the 16S rRNA gene metabarcoding, metabolomics, and multi-omics analyses can be found at https://github.com/BenoitPAIX/Rariphotic_sponge_microbiomes. Sequences for sponge and associated fauna barcoding were deposited and are publicly available in GenBank under the accession numbers XXX and XXX, for 16S and 28S rRNA genes respectively (*note: submissions SUB15025070 and SUB15027471 are currently being processed by GenBank. Datasets were submitted on the 23rd of January 2025. The dataset is temporarily available on the “BenoitPAIX/Rariphotic_sponge_microbiomes” Github repository*).

### Competing interests

The authors declare that they have no competing interests.

## Funding

This work was funded by the NWO-VIDI with project number 16.161.301.

### Authors’ contributions

BP and NdV designed the study. BP, NdV, and NW performed the fieldwork. AT and BP performed the lab work for the 16S metabarcoding. AT, BP, and NW performed the 28S barcoding of Demospongiae samples. NdV performed the lab work for the morphological identification of the Demospongiae samples. CD performed the 16S barcoding of the Hexactinellida samples. CD and NdV performed the morphological identification of the Hexactinellida samples. AT and OE performed the lab work for the metabolomics study. BP and AT analyzed the metabarcoding results. AT, OE, and BP processed the metabolomics raw data. BP analyzed and annotated the metabolomics results. BP performed the multi-omics data integration and analyzed the results. BP and NdV wrote the first draft of the manuscript. All co-authors read, contributed to the revision, and approved the submitted version.

## Acknowledgments

A special thanks goes to Adriaan “Dutch” Schrier for giving us the opportunity to use the ‘Curasub’ submarine, and the submarine pilots Bruce Brandt, and Tico Christiaanse for making the expedition run smoothly. We also thank Jeroen van Kuijk, Alexander Swaen, Kristina Beemsterboer and Stijn Raemaekers who helped during the sampling at Curaçao. We are also grateful to Pierre-Etienne Choley for his help on the nanopore data processing, and Sam Afoullouss for the confirmation of aphrocallistin identification based on the MS/MS fragmentation. We also acknowledge Charles Fransen and Werner de Gier for the identification of the crustacean specimens.

## Notes

### Competing Interest Statement

The authors have declared no competing interest.

