## Supplementary information for "Adaptive strategies of Caribbean sponge holobionts beyond the mesophotic zone"

### Supplementary material and methods

#### Supplementary information on the metabolomics data processing

The parameters used for the data processing with Metaboscape through the T-ReX 3D processing workflow are:

| **Parameter** | **Positive ion mode** |
|---|---|
| **Peak Detection** | |
| Intensity threshold (counts) | 6000 |
| Minimum Peak Length (spectra) | 11 |
| Feature Signal | Intensity |
| **Calibration** | |
| Mass recalibration *RT* | auto-detect |
| Mass recalibration *List* | Na Formate pos |
| **Recursive Feature Extraction** | |
| Minimum Peak Length *recursive* (spectra) | 7 |
| Minimum # Features for Extraction (analysis) | 1 |
| Bucket Filter | |
| Presence of features in minimum # of analysis (analysis) | 1 |
| **MS/MS Import Configuration** | |
| Perform MS/MS import | Yes |
| MS/MS import method | Maxsum* |
| Group by collision energy | Yes |
| **Ranges** | |
| Mass Range *Start* (m/z) | 100 |
| Mass Range *End* (m/z) | 1650 |
| Keep isotope pattern information from all | Yes |
| **Ion Deconvolution** | |
| EIC Correlation | 0.8 |
| Primary Ion | $[M+H]^+$ |
| Seed Ions | $[M+Na]^+$; $[M+K]^+$ |
| Common Ions | $[M-H2O+H]^+$ |

* In case multiple MS/MS spectra exist for the same precursor, then the intensities of each MS/MS spectrum are summed and the one with the highest sum is selected.

#### Supplementary information on the building of the FBMN within GNPS

The precursor ion mass tolerance was set to 0.05 Da and the MS/MS fragment ion tolerance to 0.05 Da. A feature-based molecular network (FBMN) was then created where edges were filtered to have a cosine score above 0.70 and more than 5 matched peaks. Further, edges between two nodes were kept in the network if and only if each of the nodes appeared in each others respective top 10 most similar nodes. Finally, the maximum size of a molecular family was set to 100, and the lowest scoring edges were removed from molecular families until the molecular family size was below this threshold. The spectra in the network were then searched against GNPS spectral libraries (Wang et al., 2016; Horai et al., 2010). The library spectra were filtered in the same manner as the input data. All matches kept between network spectra and library spectra were required to have a score above 0.7 and at least 5 matched peaks.

### Supplementary tables

**Table S1. Summary of the sample list.** The color corresponds to the color codes used for all figures. * indicates sponge species that could not be collected at least in triplicate in the same photic zone.

| Sample name | Class | Species | Photic zone | Depth (m) | Sampling date |
|---|---|---|---|---|---|
| Ps1.1 | Demospongiae | *Petrosia* sp. 1 * | Mesophotic | 89 | 23.04.2022 |
| Ne.1 | Demospongiae | *Neopetrosia eurystomata* | Mesophotic | 94 | 23.04.2022 |
| Sz.1 | Demospongiae | *Svenzea zeai* | Mesophotic | 95 | 23.04.2022 |
| Sz.3 | Demospongiae | *Svenzea zeai* | Mesophotic | 95 | 23.04.2022 |
| Ck.1 | Demospongiae | *Cinachyrella kuekenthali* * | Mesophotic | 100 | 23.04.2022 |
| Sz.2 | Demospongiae | *Svenzea zeai* | Mesophotic | 100 | 23.04.2022 |
| Gc.1 | Demospongiae | *Geodia* aff. *curacaoensis* | Mesophotic | 105 | 23.04.2022 |
| Ne.3 | Demospongiae | *Neopetrosia eurystomata* | Mesophotic | 105 | 23.04.2022 |
| Ne.4 | Demospongiae | *Neopetrosia eurystomata* | Mesophotic | 105 | 23.04.2022 |
| Pw.1 | Demospongiae | *Petrosia* aff. *weinbergi* * | Mesophotic | 106 | 23.04.2022 |
| Ne.2 | Demospongiae | *Neopetrosia eurystomata* | Mesophotic | 107 | 23.04.2022 |
| Gs1.1 | Demospongiae | *Geodia* sp. 1 * | Mesophotic | 121 | 23.04.2022 |
| Ah.2 | Demospongiae | *Aciculites higginsii* | Upper rariphotic | 127 | 23.04.2022 |
| Ah.3 | Demospongiae | *Aciculites higginsii* | Upper rariphotic | 127 | 23.04.2022 |
| Ah.1 | Demospongiae | *Aciculites higginsii* | Upper rariphotic | 129 | 23.04.2022 |
| Ps2.1 | Demospongiae | *Petrosia* sp. 2 * | Upper rariphotic | 129 | 23.04.2022 |
| Gi.1 | Demospongiae | *Gastrophanella implexa* * | Upper rariphotic | 151 | 23.04.2022 |
| B.m.1 | Demospongiae | *Biemna microacanthosigma* * | Lower rariphotic | 258 | 23.04.2022 |
| Pm.1 | Demospongiae | *Penares mastoideus* * | Lower rariphotic | 262 | 23.04.2022 |
| Gm.1 | Demospongiae | *Geodia* cf. *megastrella* * | Lower rariphotic | 266 | 23.04.2022 |
| Cl.2 | Demospongiae | *Calthropella lithistina* * | Lower rariphotic | 267 | 23.04.2022 |
| Gs2.1 | Demospongiae | *Geodia* sp. 2 * | Lower rariphotic | 267 | 23.04.2022 |
| To.1 | Demospongiae | *Topsentia ophiraphidites* * | Lower rariphotic | 295 | 22.04.2022 |
| Cl.1 | Demospongiae | *Calthropella lithistina* * | Lower rariphotic | 300 | 22.04.2022 |
| Dp.8 | Hexactinellida | *Dactylocalyx pumiceus* | Upper rariphotic | 132 | 30.03.2023 |
| Dp.9 | Hexactinellida | *Dactylocalyx pumiceus* | Upper rariphotic | 144 | 30.03.2023 |
| Dp.5 | Hexactinellida | *Dactylocalyx pumiceus* | Upper rariphotic | 145 | 30.03.2023 |
| Dp.6 | Hexactinellida | *Dactylocalyx pumiceus* | Upper rariphotic | 145 | 30.03.2023 |
| Dp.7 | Hexactinellida | *Dactylocalyx pumiceus* | Upper rariphotic | 145 | 30.03.2023 |
| Dp.3 | Hexactinellida | *Dactylocalyx pumiceus* | Upper rariphotic | 174 | 23.04.2022 |
| Dp.4 | Hexactinellida | *Dactylocalyx pumiceus* | Upper rariphotic | 174 | 23.04.2022 |
| Vl.6 | Hexactinellida | *Verrucocoeloidea liberatorii* | Upper rariphotic | 176 | 30.03.2023 |
| Vl.7 | Hexactinellida | *Verrucocoeloidea liberatorii* | Upper rariphotic | 176 | 30.03.2023 |
| Vl.8 | Hexactinellida | *Verrucocoeloidea liberatorii* | Upper rariphotic | 176 | 30.03.2023 |
| Ls.4 | Hexactinellida | *Lefroyella* sp. * | Upper rariphotic | 194 | 30.03.2023 |
| Cp.5 | Hexactinellida | *Conorete pourtalesi* | Lower rariphotic | 238 | 30.03.2023 |
| Cp.6 | Hexactinellida | *Conorete pourtalesi* | Lower rariphotic | 238 | 30.03.2023 |
| Cp.7 | Hexactinellida | *Conorete pourtalesi* | Lower rariphotic | 238 | 30.03.2023 |
| Hx.2 | Hexactinellida | *Hexactinella* sp. | Lower rariphotic | 238 | 30.03.2023 |
| Hx.3 | Hexactinellida | *Hexactinella* sp. | Lower rariphotic | 238 | 30.03.2023 |
| Cp.2 | Hexactinellida | *Conorete pourtalesi* | Lower rariphotic | 262 | 23.04.2022 |

| | | | | | |
|---|---|---|---|---|---|
| Cp.4 | Hexactinellida | *Conorete pourtalesi* | Lower rariphotic | 262 | 23.04.2022 |
| Dp.2 | Hexactinellida | *Dactylocalyx pumiceus* | Lower rariphotic | 262 | 23.04.2022 |
| Cp.8 | Hexactinellida | *Conorete pourtalesi* | Lower rariphotic | 267 | 30.10.2018 |
| Cp.3 | Hexactinellida | *Conorete pourtalesi* | Lower rariphotic | 268 | 23.04.2022 |
| Dp.1 | Hexactinellida | *Dactylocalyx pumiceus* | Lower rariphotic | 268 | 23.04.2022 |
| Vl.1 | Hexactinellida | *Verrucocoeloidea liberatorii* | Lower rariphotic | 268 | 23.04.2022 |
| Vl.2 | Hexactinellida | *Verrucocoeloidea liberatorii* | Lower rariphotic | 268 | 23.04.2022 |
| Vl.3 | Hexactinellida | *Verrucocoeloidea liberatorii* | Lower rariphotic | 268 | 23.04.2022 |
| Vl.4 | Hexactinellida | *Verrucocoeloidea liberatorii* | Lower rariphotic | 268 | 23.04.2022 |
| Vl.5 | Hexactinellida | *Verrucocoeloidea liberatorii* | Lower rariphotic | 268 | 23.04.2022 |
| Ls.1 | Hexactinellida | *Lefroyella* sp. | Lower rariphotic | 282 | 22.04.2022 |
| Ht.1 | Hexactinellida | *Heterotella* sp. nov. | Lower rariphotic | 288 | 22.04.2022 |
| Ht.2 | Hexactinellida | *Heterotella* sp. nov. | Lower rariphotic | 288 | 22.04.2022 |
| Cp.1 | Hexactinellida | *Conorete pourtalesi* | Lower rariphotic | 298 | 22.04.2022 |
| Hx.1 | Hexactinellida | *Hexactinella* sp. | Lower rariphotic | 301 | 22.04.2022 |
| Ls.2 | Hexactinellida | *Lefroyella* sp. | Lower rariphotic | 301 | 22.04.2022 |
| Ls.3 | Hexactinellida | *Lefroyella* sp. | Lower rariphotic | 301 | 22.04.2022 |
| Ms.1 | Hexactinellida | *Myliusia* sp. nov. * | Lower rariphotic | 302 | 22.04.2022 |
| Ht.3 | Hexactinellida | *Heterotella* sp. nov. | Lower rariphotic | 305 | 30.10.2018 |
| Sponge associated fauna | | | | | |
| NA | Anthozoa | *Vitrumanthus schrieri* | Lower rariphotic | 268 | 23.04.2022 |
| NA | Malacostraca | *Eiconaxius caribbaeus* | Lower rariphotic | 262 | 23.04.2022 |

**Table S2. Results of the two-way PERMANOVA tests conducted with the sponge class and the photic zone factors, for both metabarcoding and metabolomics datasets.** D.f., F, $R^2$ and *p* correspond to degrees of freedom, F ratio, coefficient of determination, and *p*-value, respectively

| Dataset | Factor tested | D.f. | Sum of squares | $R^2$ | F | *p* |
|---|---|---|---|---|---|---|
| **Meta-barcoding** | Sponge class | 1 | 2.25974339 | 0.08591288 | 5.8653859 | **0.001** |
| | Photic zone | 2 | 2.40547206 | 0.09145332 | 3.12181949 | **0.001** |
| | Sponge class * Photic zone | 1 | 1.2183296 | 0.04631951 | 3.16229412 | **0.001** |
| | Residual | 53 | 20.419185 | 0.77631428 | NA | NA |
| | Total | 57 | 26.3027301 | 1 | NA | NA |
| **Metabo-lomics** | Sponge class | 1 | 3872.228 | 0.070284 | 4.451549 | **0.001** |
| | Photic zone | 2 | 2300.252 | 0.041752 | 1.322195 | **0.006** |
| | Sponge class * Photic_zone | 1 | 1078.9 | 0.019583 | 1.240313 | **0.05** |
| | Residual | 55 | 47842.35 | 0.868381 | NA | NA |
| | Total | 59 | 55093.73 | 1 | NA | NA |

**Table S3. Results of the multivariate pairwise adonis test comparing five groups of sponge samples gathered according to their sponge class and photic zone, for both metabarcoding and metabolomics datasets.** M, UR, and LR correspond to the mesophotic, upper rariphotic, and lower rariphotic zones, respectively. F, $R^{2,}$ and *p* correspond to the F ratio, coefficient of determination, and *p*-value, respectively

| Dataset | Pairwise comparison | Sum of squares | F | $R^2$ | *p* |
|---|---|---|---|---|---|
| Meta-barcoding | Hexactinellida LR *vs* Demospongiae M | 0.0130 | 6.241 | 0.155 | **0.003** |
| | Hexactinellida LR *vs* Demospongiae LR | 0.0120 | 5.369 | 0.156 | **0.006** |
| | Hexactinellida LR *vs* Hexactinellida UR | 0.0074 | 3.006 | 0.081 | **0.052** |
| | Hexactinellida LR *vs* Demospongiae UR | 0.0090 | 3.859 | 0.121 | **0.019** |
| | Demospongiae M *vs* Demospongiae LR | 0.0043 | 7.748 | 0.341 | **0.001** |
| | Demospongiae M *vs* Hexactinellida UR | 0.0098 | 7.280 | 0.267 | **0.001** |
| | Demospongiae M *vs* Demospongiae UR | 0.0031 | 5.036 | 0.265 | **0.005** |
| | Demospongiae LR *vs* Hexactinellida UR | 0.0096 | 6.979 | 0.318 | **0.001** |
| | Demospongiae LR *vs* Demospongiae UR | 0.0032 | 12.343 | 0.578 | **0.003** |
| | Hexactinellida UR *vs* Demospongiae UR | 0.0074 | 4.956 | 0.261 | **0.022** |
| Metabo-lomics | Hexactinellida LR *vs* Demospongiae M | 2583.95 | 2.901 | 0.077 | **0.001** |
| | Hexactinellida LR *vs* Demospongiae LR | 2059.88 | 2.348 | 0.073 | **0.001** |
| | Hexactinellida LR *vs* Hexactinellida UR | 974.32 | 1.120 | 0.032 | **0.14** |
| | Hexactinellida LR *vs* Demospongiae UR | 2092.46 | 2.342 | 0.077 | **0.001** |
| | Demospongiae LR *vs* Demospongiae M | 1212.10 | 1.405 | 0.076 | **0.008** |
| | Demospongiae LR *vs* Demospongiae UR | 1083.63 | 1.274 | 0.113 | **0.023** |
| | Demospongiae LR *vs* Hexactinellida UR | 1868.60 | 2.287 | 0.125 | **0.001** |
| | Demospongiae M *vs* Demospongiae UR | 1275.44 | 1.432 | 0.087 | **0.009** |
| | Demospongiae M *vs* Hexactinellida UR | 2294.26 | 2.689 | 0.114 | **0.001** |
| | Demospongiae UR *vs* Hexactinellida UR | 1974.53 | 2.350 | 0.144 | **0.001** |

**Table S4. Results of the nested PERMANOVA tests conducted with the "sponge genus" factor nested within each "sponge class" factor, for both metabarcoding and metabolomics datasets.** D.f., F, $R^2$ and *p* correspond to degrees of freedom, F ratio, coefficient of determination, and *p*-value, respectively

| Dataset | Factor tested | D.f. | Sum of squares | $R^2$ | F | *p* |
|---|---|---|---|---|---|---|
| Meta-barcoding | Sponge class | 1 | 2.259743 | 0.085913 | 10.82349 | **0.001** |
| | Sponge class / Sponge genus | 17 | 15.90052 | 0.60452 | 4.479924 | **0.001** |
| | Residual | 39 | 8.142471 | 0.309568 | NA | NA |
| | Total | 57 | 26.30273 | 1 | NA | NA |
| Metabo-lomics | Class_host | 1 | 3872.228 | 0.070 | 4.967 | **0.001** |
| | Sponge class / Sponge genus | 17 | 19260.480 | 0.350 | 1.453 | **0.001** |
| | Residual | 41 | 31961.019 | 0.580 | NA | NA |
| | Total | 59 | 55093.727 | 1.000 | NA | NA |

**Table S5. Summarized results of the significant multivariate pairwise adonis comparisons within sponge genus, for both metabarcoding and metabolomics datasets.** F, $R^{2,}$ and *p* correspond to the F ratio, coefficient of determination, and *p*-value, respectively.

| Dataset | Pairwise comparison | Sum of squares | F | $R^2$ | *p* |
|---|---|---|---|---|---|
| Metabar-coding | *Lefroyella vs Neopetrosia* | 0.00523906 | 41.4445051 | 0.89234464 | 0.03 |
| | *Lefroyella vs Verrucocoeloidea* | 0.01726565 | 134.439986 | 0.93076709 | 0.001 |
| | *Lefroyella vs Hexactinella* | 0.0018457 | 8.62206806 | 0.63294854 | 0.032 |
| | *Lefroyella vs Dactylocalyx* | 0.00381299 | 8.06953216 | 0.42316361 | 0.001 |
| | *Lefroyella vs Conorete* | 0.0151288 | 82.3261723 | 0.89168835 | 0.005 |
| | *Lefroyella vs Petrosia* | 0.00255957 | 8.21184424 | 0.6215517 | 0.026 |
| | *Lefroyella vs Geodia* | 0.00480402 | 12.8215718 | 0.71944113 | 0.019 |
| | *Lefroyella vs Svenzea* | 0.00309843 | 9.20254244 | 0.64795036 | 0.03 |
| | *Lefroyella vs Aciculites* | 0.00438964 | 35.503539 | 0.876554 | 0.032 |
| | *Lefroyella vs Heterotella* | 0.00099723 | 4.81729681 | 0.49069483 | 0.038 |
| | *Neopetrosia vs Verrucocoeloidea* | 0.01915322 | 200.189898 | 0.95697689 | 0.007 |
| | *Neopetrosia vs Dactylocalyx* | 0.00751745 | 15.7449931 | 0.61157496 | 0.003 |
| | *Neopetrosia vs Conorete* | 0.01758884 | 111.913674 | 0.92556673 | 0.006 |
| | *Calthropella vs Verrucocoeloidea* | 0.0128796 | 124.850423 | 0.9397819 | 0.019 |
| | *Calthropella vs Dactylocalyx* | 0.00467829 | 8.88526002 | 0.49679233 | 0.017 |
| | *Calthropella vs Conorete* | 0.01181092 | 68.534448 | 0.89547191 | 0.018 |
| | *Verrucocoeloidea vs Hexactinella* | 0.01176412 | 81.484416 | 0.90053536 | 0.008 |
| | *Verrucocoeloidea vs Dactylocalyx* | 0.03102167 | 85.7471905 | 0.85111247 | 0.001 |
| | *Verrucocoeloidea vs Conorete* | 0.05495612 | 372.279926 | 0.96375685 | 0.001 |
| | *Verrucocoeloidea vs Petrosia* | 0.01388056 | 69.8887473 | 0.88591529 | 0.01 |
| | *Verrucocoeloidea vs Geodia* | 0.01884016 | 80.6500849 | 0.89960969 | 0.005 |
| | *Verrucocoeloidea vs Svenzea* | 0.01409991 | 66.3529963 | 0.88056215 | 0.011 |
| | *Verrucocoeloidea vs Aciculites* | 0.0173875 | 184.707594 | 0.95353822 | 0.006 |
| | *Verrucocoeloidea vs Heterotella* | 0.01113463 | 79.2769507 | 0.89804813 | 0.009 |
| | *Hexactinella vs Dactylocalyx* | 0.00294304 | 5.64582675 | 0.36085193 | 0.003 |
| | *Hexactinella vs Conorete* | 0.01122939 | 54.5482095 | 0.85837524 | 0.006 |
| | *Dactylocalyx vs Conorete* | 0.02736767 | 68.6467152 | 0.82067437 | 0.001 |
| | *Dactylocalyx vs Petrosia* | 0.00359733 | 6.31009759 | 0.38688288 | 0.007 |
| | *Dactylocalyx vs Geodia* | 0.006976 | 11.5960222 | 0.53695176 | 0.004 |
| | *Dactylocalyx vs Svenzea* | 0.00446347 | 7.66141307 | 0.43379389 | 0.008 |
| | *Dactylocalyx vs Aciculites* | 0.00629095 | 13.2144952 | 0.56923466 | 0.009 |
| | *Dactylocalyx vs Heterotella* | 0.00190949 | 3.68805768 | 0.26943616 | 0.012 |
| | *Conorete vs Petrosia* | 0.01248329 | 47.9944898 | 0.84209 | 0.002 |
| | *Conorete vs Geodia* | 0.01723654 | 58.4105863 | 0.86648981 | 0.006 |
| | *Conorete vs Svenzea* | 0.01148638 | 41.9229762 | 0.82326249 | 0.005 |
| | *Conorete vs Aciculites* | 0.01597357 | 102.641788 | 0.91938502 | 0.012 |
| | *Conorete vs Heterotella* | 0.00959747 | 47.5259862 | 0.84078119 | 0.005 |
| Metabo-lomics | *Conorete vs Lefroyella* | 1401.263345 | 1.689045381 | 0.144498146 | 0.003 |
| | *Conorete vs Hexactinella* | 1113.436064 | 1.314648936 | 0.12745455 | 0.049 |
| | *Conorete vs Calthropella* | 1600.656439 | 1.889794356 | 0.191085303 | 0.017 |
| | *Conorete vs Svenzea* | 1612.116222 | 1.953908422 | 0.178375457 | 0.009 |
| | *Conorete vs Neopetrosia* | 2190.840979 | 2.628688083 | 0.208152111 | 0.004 |
| | *Conorete vs Petrosia* | 1665.989698 | 1.896363957 | 0.1740364 | 0.005 |
| | *Conorete vs Aciculites* | 2044.543599 | 2.355695111 | 0.207446139 | 0.005 |
| | *Conorete vs Geodia* | 1835.762546 | 2.21888216 | 0.18159453 | 0.001 |
| | *Conorete vs Dactylocalyx* | 1758.044958 | 2.191615014 | 0.127481625 | 0.001 |
| | *Conorete vs Verrucocoeloidea* | 1332.986913 | 1.611870356 | 0.103246461 | 0.008 |
| | *Heterotella vs Neopetrosia* | 1608.428747 | 1.944174546 | 0.279972016 | 0.042 |
| | *Heterotella vs Geodia* | 1142.557535 | 1.401722198 | 0.218960173 | 0.028 |

| | | | | | |
|---|---|---|---|---|---|
| | *Heterotella vs Dactylocalyx* | 1055.502635 | 1.34721289 | 0.118726325 | 0.038 |
| | *Heterotella vs Verrucocoeloidea* | 1341.434393 | 1.635912643 | 0.153810275 | 0.012 |
| | *Lefroyella vs Svenzea* | 1127.614706 | 1.586429832 | 0.240863392 | 0.026 |
| | *Lefroyella vs Neopetrosia* | 1673.36145 | 2.249828992 | 0.272712197 | 0.03 |
| | *Lefroyella vs Petrosia* | 1359.286813 | 1.684392689 | 0.251988889 | 0.033 |
| | *Lefroyella vs Aciculites* | 1665.753705 | 2.114157155 | 0.297176055 | 0.037 |
| | *Lefroyella vs Geodia* | 1231.778602 | 1.679067798 | 0.21865516 | 0.027 |
| | *Lefroyella vs Dactylocalyx* | 1211.856755 | 1.633467169 | 0.129296823 | 0.004 |
| | *Lefroyella vs Verrucocoeloidea* | 1490.199462 | 1.933810661 | 0.162044691 | 0.003 |
| | *Hexactinella vs Neopetrosia* | 1528.282679 | 2.016756931 | 0.287420093 | 0.023 |
| | *Hexactinella vs Geodia* | 1279.116247 | 1.715563797 | 0.255460874 | 0.026 |
| | *Hexactinella vs Dactylocalyx* | 1031.760591 | 1.378043633 | 0.121114286 | 0.008 |
| | *Hexactinella vs Verrucocoeloidea* | 1092.403853 | 1.398058149 | 0.134453773 | 0.008 |
| | *Calthropella vs Dactylocalyx* | 1540.655338 | 2.088038631 | 0.188314516 | 0.016 |
| | *Calthropella vs Verrucocoeloidea* | 1700.631084 | 2.199384729 | 0.215638961 | 0.029 |
| | *Svenzea vs Neopetrosia* | 1321.780668 | 1.83984181 | 0.268988942 | 0.024 |
| | *Svenzea vs Geodia* | 1135.189711 | 1.607407697 | 0.243273576 | 0.034 |
| | *Svenzea vs Dactylocalyx* | 1697.840875 | 2.328909319 | 0.188898244 | 0.011 |
| | *Svenzea vs Verrucocoeloidea* | 1826.588713 | 2.404990743 | 0.210871784 | 0.005 |
| | *Neopetrosia vs Petrosia* | 1350.887486 | 1.658298862 | 0.249057439 | 0.029 |
| | *Neopetrosia vs Aciculites* | 1800.607826 | 2.263384999 | 0.311615727 | 0.026 |
| | *Neopetrosia vs Geodia* | 1661.47933 | 2.245334557 | 0.272315761 | 0.03 |
| | *Neopetrosia vs Dactylocalyx* | 2254.248316 | 3.024367282 | 0.21565089 | 0.002 |
| | *Neopetrosia vs Verrucocoeloidea* | 2322.923449 | 2.999568655 | 0.230743707 | 0.002 |
| | *Petrosia vs Geodia* | 1092.317003 | 1.36126903 | 0.213993312 | 0.025 |
| | *Petrosia vs Dactylocalyx* | 1742.177094 | 2.241811548 | 0.183127435 | 0.003 |
| | *Petrosia vs Verrucocoeloidea* | 1891.448956 | 2.326663971 | 0.205414761 | 0.005 |
| | *Aciculites vs Geodia* | 1226.260616 | 1.565424475 | 0.238434618 | 0.03 |
| | *Aciculites vs Dactylocalyx* | 2099.309444 | 2.734947617 | 0.214759236 | 0.012 |
| | *Aciculites vs Verrucocoeloidea* | 2257.174676 | 2.813232913 | 0.238142508 | 0.008 |
| | *Geodia vs Dactylocalyx* | 1560.413681 | 2.109186183 | 0.160893754 | 0.001 |
| | *Geodia vs Verrucocoeloidea* | 1884.724231 | 2.453043687 | 0.196983464 | 0.006 |
| | *Dactylocalyx vs Verrucocoeloidea* | 1960.099173 | 2.569527594 | 0.1462491 | 0.001 |

**Table S6. List of biomarker metabolites (VIP score > 2) annotated by LC-HRMS/MS and involved in the discriminations between the two sponge classes (Demospongiae *vs* Hexactinellida).** *P* values correspond to the result of the T-test using the "sponge class" factor. Color codes correspond to mean normalized concentrations. Annotation confidence levels are determined according to Schymanski et al., 2014. [a] Constructor statistical match factor (comparison of theoretical and experimental isotopic patterns); [b] Abbreviations: PC: phosphatidylcholine; PE: phosphatidylethanolamine; PI: phosphatidylinositol; n.i.: not identified; n.f.: not fragmented.

| VIP number | VIP score Comp. 1 | *m/z* | RT (s) | Molecular formula | Adduct | Mass err (ppm) | mσ [a] | Annotation confidence | Putative annotation [b] | MSMS fragments | T-test p-values | Demo-spongiae | Hexacti-nellida |
|---|---|---|---|---|---|---|---|---|---|---|---|---|---|
| VIP 01 | 4.08 | 510.3521 | 960 | $C_{25}H_{52}NO_7P$ | $[M+H]^+$ | 6.5 | 14.3 | 2a | *Lyso* -PC(C17:0) | 184.0721 $[C_5H_{15}NO_4P]^+$ ; 258.1065 $[C_8H_{21}NO_6P]^+$ | < 2.2e-16 | | |
| VIP 02 | 4.03 | 261.1412 | 276 | $C_{11}H_{20}N_2O_5$ | $[M+H]^+$ | 12.8 ? | 11.3 | 3 | Glutamylisoleucine ? | 130.0478 $[C_5H_8NO_3]^+$ | < 2.2e-16 | | |
| VIP 03 | 3.95 | 544.3351 | 900 | $C_{28}H_{50}NO_7P$ | $[M+H]^+$ | 2.2 | 73 | 2a | *Lyso* -PC(C20:4) | 526.3283 $[C_{28}H_{49}NO_6P]^+$ ; 258.1132 $[C_8H_{21}NO_6P]^+$ ; 184.0724 $[C_{15}H_{15}NO_5P]^+$ | < 2.2e-16 | | |
| VIP 04 | 3.83 | 524.3677 | 1068 | $C_{26}H_{54}NO_7P$ | $[M+H]^+$ | 6.5 | n.a. | 2a | *Lyso* -PC(C18:0) | 506.3622 $[C_{26}H_{53}NO_6P]^+$ ; 258.0952 $[C_8H_{21}NO_6P]^+$ ; 184.0720 $[C_{15}H_{15}NO_5P]^+$ | < 2.2e-16 | | |
| VIP 05 | 3.52 | 264.157 | 126 | $C_{10}H_{18}N_2O_5$ | $[M+NH_4]^+$ | 2.4 | 0.63 | 3 | Glutamylvaline ? | 162.0839 $[C_5H_{12}N_3O_3]+$ ; 170.0893 $[C_7H_{12}N_3O_2]+$ | 3.31E-12 | | |
| VIP 08 | 3.42 | 190.0455 | 66 | $C_8H_9NO_3$ | $[M+Na]^+$ | 10.4 | 11.8 | 4 | Hydroxyphenylglycine ? | n.f. | 5.617E-13 | | |
| VIP 09 | 3.39 | 168.0647 | 72 | $C_8H_9NO_3$ | $[M+H]^+$ | 4.6 | 13.8 | 4 | Hydroxyphenylglycine ? | n.f. | 5.617E-13 | | |
| VIP 10 | 3.36 | 496.3727 | 1026 | $C_{25}H_{54}NO_6P$ | $[M+H]^+$ | 8.2 | n.a. | 2a | *Lyso* -PC(O-C17:0) | 204.0864 ; 184.0724 $[C_{15}H_{15}NO_4P]^+$ | 5.669E-10 | | |
| VIP 11 | 3.32 | 542.3205 | 846 | $C_{28}H_{48}NO_7P$ | $[M+H]^+$ | 6.7 | n.a. | 2a | *Lyso* -PC(C20:5) | 507.4401 ; 235.1931 ; 198.0814 ; 184.0736 $[C_{15}H_{15}NO_5P]^+$ | 4.457E-11 | | |
| VIP 12 | 3.06 | 241.1536 | 96 | $C_{12}H_{20}N_2O_3$ | $[M+H]^+$ | 4.3 | n.a. | 4 | Unidentified $C_{12}H_{20}N_2O_3$ | 149.1148 $[C_7H_{17}O_3]^+$ ; 181.1005 $[C_9H_{13}N_2O_2]^+$ ; 196.0969 $[C_{10}H_{14}NO_3]^+$ | 3.928E-09 | | |
| VIP 13 | 3.04 | 539.0407 / 541.0402 | 522 | $C_{20}H_{24}Br_2N_6O_2$ | $[M+H]^+$ | -0.4 | 21.1 | 2a | Aphrocallistin | 134.0602 $[C_8H_8NO]^+$ ; 148.0616 $[C_6H_6N_5]^+$ ; 163.0865 $[C_7H9N5]^+$ ; 176.0930 $[C_8H_{10}N_5]^+$ ; 190.1080 $[C_9H_{12}N_5]^+$ ; 374.0587 $[C_{15}H_{21}BrN_5O]^+$ ; 376.08 $[C_{16}H_{19}BrN_5O]^+$ | 0.000004345 | | |
| VIP 14 | 3.04 | 508.3408 | 900 | $C_{25}H_{50}NO_7P$ | $[M+H]^+$ | -2.1 | n.a. | 2a | *Lyso* -PC(C17:1) | 184.0731 $[C_{15}H_{15}NO_5P]^+$ | 4.791E-08 | | |
| VIP 15 | 2.92 | 231.0819 | 90 | $C_9H_{14}N_2O_3S$ | $[M+H]^+$ | -9.3 | 33.1 | 4 | Methionylalanine derivative ? | 146.0811 $[C_6H_{12}NO_3]^+$ | 5.626E-08 | | |
| VIP 16 | 2.70 | 510.3872 | 1068 | $C_{26}H_{56}NO_6P$ | $[M+H]^+$ | 6.5 | n.a. | 2a | *Lyso* -PC(O-C18:0) | 184.0725 $[C_5H_{15}NO_4P]^+$ ; 240.1040 $[C_8H_{19}NO_5P]^+$ ; 258.1134 $[C_8H_{21}NO_6P]^+$ | 8.208E-07 | | |
| VIP 17 | 2.68 | 538.3838 | 1122 | $C_{27}H_{56}NO_7P$ | $[M+H]^+$ | 5.4 | 64.4 | 2a | *Lyso* -PC(C19:0) | 184.0721 $[C_5H_{15}NO_4P]+$ ; 258.1055 $[C_8H_{21}NO_6P]+$ ; 355.3181 $[C_{22}H_{43}O_3]+$ ; 520.3720 $[C_{27}H_{55}NO_6P]+$ | 8.819E-09 | | |
| VIP 18 | 2.63 | 198.0848 | 72 | $C_8H_{11}N_3O_3$ | $[M+H]^+$ | 12.7 | 24.8 | 3 | Methylated aminoacid derivative | 184.0667 $[M-CH_2+H]+$ $[C_7H_{10}N_3O_3]^+$ ; 139.0127 $[C_5H_3N_2O_3]^+$ | 3.114E-07 | | |
| VIP 19 | 2.58 | 597.3366 | 1134 | $C_{28}H_{53}O_{11}P$ | $[M+H-H_2O]^+$ | 5.4 | 63.1 | 3 | *Lyso* -PI(C19:0) ? | 155.0107 $[C_3H_8O_5P]^+$ | 0.00001875 | | |
| VIP 20 | 2.44 | 550.3618 | 1092 | $C_{28}H_{56}NO_7P$ | $[M+H]^+$ | 6.8 | 50.7 | 2a | *Lyso* -PC(C20:1) | 532.3716 $[C_{28}H_{55}NO_6P]^+$ $[M-H_2O+H]^+$ ; 184.07 $[C_{15}H_{15}NO_5P]^+$ | 0.000004104 | | |
| VIP 21 | 2.43 | 187.1075 | 66 | $C_8H_{14}N_2O_3$ | $[M+H]^+$ | 6.3 | 4.7 | 3 | Alanylproline | 141.050 $[C_7H_{13}N_2O]^+$ | 2.076E-08 | | |
| VIP 22 | 2.41 | 271.0231 | 522 | $C_{20}H_{24}Br_2N_6O_2$ | 2[M+H]+ | -0.4 | 21.1 | 2a | Aphrocallistin fragment | 134.0602 $[C_8H_8NO]^+$ ; 163.0865 $[C_9H_{11}N_2O]^+$ ; 190.1079 $[C_8H_{16}NO_4]^+$ ; 197.9659 $[C_6H_5BrN_3]^+$ ? | 0.0001999 | | |
| VIP 23 | 2.41 | 583.3221 | 1074 | $C_{27}H_{53}O_{12}P$ ? | $[M+H-H_2O]^+$ | 3.6 | n.a. | 3 | *Lyso* -PI(C18:0) ? | 155.0097 $[C_3H_8O_5P]^+$ | 0.000001701 | | |
| VIP 24 | 2.41 | 502.2896 | 864 | $C_{25}H_{44}NO_7P$ | $[M+H]^+$ | 6.3 | 32.2 | 2a | *Lyso* -PE(C20:4) | 361.2854 $[M-141.0189]^+$ $[M-C_2H_8NO_4P]^+$ $[C_{23}H_{37}O_3]^+$ | 0.00002457 | | |
| VIP 25 | 2.39 | 569.0453 | 504 | $C_{21}H_{26}Br_2N_6O_3$ | $[M+H]^+$ | 9.3 | 23.7 | 2a | Aphrocallistin derivative | 527.0654 $[C_{19}H_{25}Br_2N_6O_2]^+$ ; 220.1176 $[C_{10}H_{14}N_5O]^+$ ; 206.0986 $[C_9H_{12}N_5O]^+$ ; 193.0951 $[C_{10}H_{13}N_2O_2]^+$ ; 178.1175 $[C_8H_{12}N_5]^+$ ; 137.0832 $[C_6H_9N_4]^+$ | 0.00009249 | | |
| VIP 26 | 2.24 | 189.103 | 108 | $C_{11}H_9NO$ | $[M+NH_4]^+$ | -4 | 41.7 | 4 | n.i. | 172.0771 $[C_{11}H_{10}NO]^+$ ; $[M+H]^+$ ; 160.0719 $[C_{10}H_{10}NO]^+$ | 1.299E-09 | | |
| VIP 27 | 2.11 | 206.0444 | 276 | $C_{10}H_7NO_4$ | $[M+H]^+$ | 1.7 | 4.6 | 2a | Xanthurenic acid | 132.0436 $[C_8H_6NO]^+$ ; 160.377 $[C_9H_6NO_2]^+$ ; 178.0484 $[C_9H_8NO_3]^+$ | 0.001329 | | |
| VIP 28 | 2.07 | 246.1685 | 324 | $C_{12}H_{23}NO_4$ | $[M+H]^+$ | 3.8 | 10 | 2b | Valerylcarnitine | 187.0989 $[C_9H_{15}O_4]^+$ ; 144.1036 $[C_7H_{14}NO_2]^+$ ; 133.0459 $[C_5H_9O_4]^+$ | 0.0001351 | | |

| | | |
|---|---|---|
| Color code for normalized concentrations | -1.2 | 1.2 |

**Table S7. List of biomarker metabolites (VIP score > 3) annotated by LC-HRMS/MS and involved in the photic zone discriminations for the Demospongiae samples. M, UR, and LR corresponds to the mesophotic, upper rariphotic, and lower rariphotic zones, respectively).** *P* values correspond to the result of the one-way ANOVA test using the "photic zone" factor. Color codes correspond to mean normalized concentrations. Annotation confidence levels are determined according to Schymanski et al., 2014.[a] Constructor statistical match factor (comparison of theoretical and experimental isotopic patterns). [b] Abbreviations: PC: phosphatidylcholine; PAF: platelet activating factor; n.i.: not identified; n.f.: not fragmented. [c] Compound previously identified in Table S6.

| VIP number | VIP score Comp. 1 | *m/z* | RT (s) | Molecular formula | Adduct | Mass err (ppm) | mσ [a] | Annotation confidence | Putative annotation [b] | MSMS fragments | ANOVA *p*-values | M | UR | LR |
|---|---|---|---|---|---|---|---|---|---|---|---|---|---|---|
| VIP 16 [c] | 5.0305 | 510.3872 | 1068 | $C_{26}H_{56}NO_6P$ | $[M+H]^+$ | 6.5 | n.a. | 2a | *Lyso*-PC(O-C18:0) (*Lyso*-PAF) | 184.0725 $[C_5H_{15}NO_4P]^+$ ; 240.1040 $[C_8H_{19}NO_5P]^+$ ; 258.1134 $[C_8H_{21}NO_6P]^+$ | 0.000203 | | | |
| VIP 30 | 4.886 | 176.1268 | 72 | $C_8H_{17}NO_3$ | $[M+H]^+$ | 7.5 | 7.2 | 2b | Rhodosamine | 174.1071 $[C_8H_{16}NO_3]^+$ ; 156.0571 $[C_7H_{10}NO_3]^+$ ; 144.0948 $[C_7H_{14}NO_2]^+$ ; 128.1012 $[C_7H_{14}NO]^+$ | 0.000427 | | | |
| VIP 31 | 4.6516 | 155.0811 | 72 | $C_7H_{10}N_2O_2$ | $[M+H]^+$ | 2.6 | 8.3 | 4 | $C_7H_{10}N_2O_2$. Cyclo-(Pro-Gly) ? | 137.0704 $[C_7H_9N_2O]$+ $[M-H_2O+H]^+$ | 9.42E-06 | | | |
| VIP 32 | 4.3934 | 174.1253 | 90 | $C_7H_{15}N_3O_2$ | $[M+H]^+$ | -9.1 | 2 | 4 | $C_7H_{15}N_3O_2$ | 157.1018 $[C_7H_{13}N_2O_2]^+$ $[M-NH_4+H]^+$ ; 156.1072 $[M-H_2O+H]^+$ ; 138.0854 | 0.00356 | | | |
| VIP 33 | 4.2348 | 522.3866 | 1218 | $C_{27}H_{56}NO_6P$ | $[M+H]^+$ | 10 | n.a. | 2a | PC(O-18:1/O-1:0) (PAF) | 184.0712 $[C_5H_{15}NO_4P]^+$ ; 304.1273 $[C_{13}H_{23}NO_5P]^+$ | 0.00508 | | | |
| VIP 34 | 4.1814 | 166.0845 | 138 | $C_9H_{11}NO_2$ | $[M+H]^+$ | 10.4 | 6.9 | 3 | $C_9H_{11}NO_2$: Phenylalanine ? | 130.0623 $[C_9H_8N]^+$ | 2.01E-05 | | | |
| VIP 35 | 3.768 | 367.1375 | 132 | $C_{19}H_{18}N_4O_4$ | $[M+H]^+$ | 6.9 | 10.8 | 4 | Unidentified alkaloid $C_{19}H_{18}N_4O_4$ | 130.0628 $[C_9H_8N]^+$ or $[C_4H_8N_3O_2]$ ; 144.0764 $[C_5H_{10}N_3O_2]^+$ ; 156.0758 $[C_6H_{10}N_3O_2]^+$ ; 170.0547 $[C_6H_8N_3O_3]^+$ ; 184.0706 $[C_7H_{10}N_3O_3]^+$ ; 202.0812 $[C_7H_{12}N_3O_4]^+$ ; 230.0740 ; 254.0770 $[C_{15}H_8N_3]^+$ ; 332.1011 $[C_{19}H_{14}N_3O_3]^+$ | 0.00609 | | | |
| VIP 36 | 3.7431 | 180.1014 | 90 | $C_{10}H_{13}NO_2$ | $[M+H]^+$ | 2.7 | 31.3 | 2b | Salsolinol | 162.0810 $[M-H_2O+H]^+$ $[C_{10}H_{12}NO]^+$ ; 136.1129 $[C_9H_{14}N]^+$ ; 134.0855 $[C_9H_{12}N]^+$ ; 121.10 $[C_9H_{13}]^+$ | 0.00393 | | | |
| VIP 37 | 3.5946 | 146.1184 | 72 | $C_7H_{15}NO_2$ | $[M+H]^+$ | -5.7 | n.a. | 4 | $C_7H_{15}NO_2$ Methyl-isoleucine ? | no fragmentation | 0.00466 | | | |
| VIP 38 | 3.427 | 137.0462 | 96 | $C_5H_4N_4O$ ? | $[M+H]^+$ | -3.3 | 22.7 | 4 | $C_5H_4N_4O$ Hypoxanthine ? | no fragmentation | 0.00119 | | | |
| VIP 39 | 3.2477 | 204.1344 | 90 | $C_8H_{18}N_3O_3$ | $[M+H]^+$ | -0.6 | 26.3 | 2b | Glycyl-Lysine (Gly-Lys) | 134.0828 $[C_5H_{12}NO_3]^+$ ; 158.1274 $[C_7H_{16}N_3O]^+$ | 0.025 | | | |
| VIP 40 | 3.2132 | 226.1189 | 72 | $C_{10}H_{17}N_3O_3$ | $[M+H-H_2O]^+$ | -1.2 | 5.5 | 2b | Glutaminyl-proline (Gln-Pro) | 208.1080 $[C_{10}H_{14}N_3O_2]^+$ ; 184.1081 $[C_8H_{14}N_3O_2]^+$ ; 182.1235 $[C_9H_{16}N_3O]^+$ ; 180.1125 $[C_9H_{14}N_3O]^+$ ; 167.0800 $[C_8H_{11}N_2O_2]^+$ ; 140.1188 $[C_7H_{14}N_3]^+$ ; 138.1037 $[C_7H_{12}N_3]^+$ | 0.0279 | | | |
| VIP 41 | 3.211 | 144.079 | 282 | $C_5H_6N_2O_2$ | $[M+NH_4]$ | -15.7 | 17.4 | 4 | $C_5H_6N_2O_2$: Thymine ? | 127.0470 $[C_5H_7N_2O_2]$+ $[M-NH_4+H]^+$ | 0.0777 | | | |
| VIP 42 | 3.0934 | 187.1075 | 66 | $C_8H_{14}N_2O_3$ | $[M+H]^+$ | 6.3 | 4.7 | 3 | Alanyl-proline (Ala-Pro) | 141.050 $[C_7H_{13}N_2O]^+$ | 0.000849 | | | |
| VIP 43 | 3.0814 | 614.3621 | 996 | $C_{37}H_{47}N_3O_5$ ? | $[M+H]^+$ | 1.2 | 90.4 | 5 | n.i. | 354.3335 $[C_{22}H_{44}NO_2]^+$ ? | 0.0335 | | | |

| Color code for normalized concentrations | | | |
|---|---|---|---|
| | -1.3 | 0 | 1.3 |

**Table S8. List of biomarker metabolites (VIP score > 3) annotation by LC-HRMS/MS and involved in the photic zone discriminations for the Hexactinellida samples. UR and LR correspond to the upper rariphotic and lower rariphotic zones, respectively).** *P* values correspond to the result of the T-test using the “photic zone” factor. Color codes correspond to mean normalized concentrations. Annotation confidence levels are determined according to Schymanski et al., 2014. [a] Constructor statistical match factor (comparison of theoretical and experimental isotopic patterns). [b] Abbreviations: PS: phosphatidylserine; PC: phosphatidylcholine; PE: phosphatidylethanolamine; PG: phosphatidylglycerol; n.i.: not identified; n.f.: not fragmented. [c] Compounds previously identified in Table S6. [d] Compound previously identified in Table S7.

| VIP number | VIP score Comp. 1 | *m/z* | RT (s) | Molecular formula | Adduct | Mass err (ppm) | mσ [a] | Annotation confidence | Putative annotation [b] | MSMS fragments | T-test p-values | UR | LR |
|---|---|---|---|---|---|---|---|---|---|---|---|---|---|
| VIP 43 | 4.969 | 482.2465 | 810 | $C_{21}H_{40}NO_9P$ | $[M+H]^+$ | 10 | 13.6 | 2a | *Lyso* -PS(C15:1) | 297.2395 $[M\text{-phosphoserine}+H]^+$ ; 242.0367 $[C_6H_{13}NO_7P]^+$ ; 223.2024 $[C_{15}H_{27}O]^+$ ; 205.1929 $[C_{15}H_{25}]^+$ ; 155.0093 $[C_3H_8O_5P]^+$ | 0.0002 | | |
| VIP 27 [c] | 4.17 | 206.0466 | 276 | $C_{10}H_7NO_4$ | $[M+H]^+$ | -8.8 | 2 | 2a | Xanthurenic acid | 132.0451 $[C_8H_6NO]^+$ ; 160.390 $[C_9H_6NO_2]^+$ ; 178.0506 $[C_9H_8NO_3]^+$ | 0.0187 | | |
| VIP 44 | 4.1524 | 570.3514 | 906 | $C_{30}H_{52}NO_7P$ | $[M+H]^+$ | 7.1 | | 2a | *Lyso* -PC(C22:5) | 184.072 $[C_5H_{15}NO_4P]^+$ | 0.0026 | | |
| VIP 45 | 3.9814 | 231.1687 | 72 | $C_{11}H_{22}N_2O_3$ | $[M+H]^+$ | 6.9 | 7 | 2b | Isoleucylvaline or Leucylvaline (Ile-Val or Leu-Val) | 130.0848 $[C_6H_{12}NO_2]^+$ ; 144.1365 $[C_8H_{18}NO]^+$; 154.0907 $[C_8H_{12}NO_2]^+$; 172.0858 $[C_8H_{14}NO_3]^+$ ; 189.1963 $[C_9H_{21}N_2O_2]^+$ | 0.0002 | | |
| VIP 46 | 3.8551 | 245.183 | 72 | $C_{12}H_{24}N_2O_3$ | $[M+H]^+$ | 12.2 | 10.2 | 2b | Leucyl-leucine (Leu-Leu) | 130.0867 $[C_6H_{12}NO_2]^+$ ; 142.1188 $[C_8H_{16}NO]^+$ ; 168.0990 $[C_9H_{14}NO_2]^+$ ;172.0975 $[C_8H_{14}NO_3]^+$; 186.1130 $[C_9H_{16}NO_3]^+$ ; 201.2020 $[C_{11}H_{25}N_2O]^+$ | 0.0003 | | |
| VIP 47 | 3.7094 | 534.3533 | 1050 | $C_{27}H_{52}NO_7P$ | $[M+H]^+$ | 4 | 20.7 | 2a | *Lyso* -PE(C22:2) | 393.3341 $[M+H\text{-}141]^+$ $[M+H\text{-}C_2H_8NO_4P]^+$ $[C_{25}H_{45}O_3]^+$ | 0.0074 | | |
| VIP 48 | 3.5729 | 794.5976 | 1170 | $C_{46}H_{84}NO_7P$ | $[M+H]^+$ | 10.4 | n.a. | 3 | PC(O-C38:5) | 184.0706 $[C_5H_{15}NO_4P]^+$ | 0.0140 | | |
| VIP 49 | 3.5465 | 188.0681 | 306 | $C_6H_9N_3O_4$ | $[M+H]^+$ | -7.9 | 25.9 | 3 | Imidazole derivative. Hydroymetronidazole ? | 170.0560 $[C_6H_8N_3O_3]^+$ ; 144.0790 $[C_5H_{10}N_3O_2]^+$ ; 142.0604 $[C_5H_8N_3O_2]^+$ | 0.0031 | | |
| VIP 50 | 3.5265 | 530.3205 | 936 | $C_{27}H_{48}NO_7P$ | $[M+H]^+$ | 2.6 | 8.9 | 2a | *Lyso* -PE(C22:4) | 389.2995 $[M+H\text{-}141]^+$ $[M+H\text{-}C_2H_8NO_4P]^+$ $[C_{25}H_{41}O_3]^+$ | 0.0124 | | |
| VIP 51 | 3.4346 | 235.165 | 66 | $C_{10}H_{22}N_2O_4$ | $[M+H]^+$ | 1 | 21.8 | 4 | Unidentified C10H22N2O4 | 176.0349 $[C_9H_6NO_3]^+$ | 0.0092 | | |
| VIP 38 [d] | 3.2128 | 204.1344 | 90 | $C_8H_{17}N_3O_3$ | $[M+H]^+$ | -0.6 | 26.3 | 2b | Glycyl-Lysine (Gly-Lys) | 134.0828 $[C_5H_{12}NO_3]^+$ ; 158.1274 $[C_7H_{16}N_3O]^+$ | 0.0605 | | |
| VIP 52 | 3.1951 | 570.3487 | 924 | $C_{30}H_{52}NO_7P$ | $[M+H]^+$ | 11.8 | 32.2 | 2a | *Lyso* -PC(C22:5) | 184.0689 $[C_5H_{15}NO_4P]^+$ | 0.0026 | | |
| VIP 53 | 3.193 | 196.0751 | 72 | $C_5H_{10}N_2O_3S$ ? | $[M+NH4]^+$ ? | -0.3 | 31 | 3 | Cysteinyl-Glycine ? | 179.0509 $[C_5H_{11}N_2O_3S]^+$ $[M\text{-}NH_4+H]^+$ | 0.0322 | | |
| VIP 54 | 3.1379 | 271.11802 | 1110 | $C_{17}H_{22}N_2O$ | $[M+H]^+$ | -8.5 | n.a. | 4 | Unidentified C17H22N2O | 138.1293 $[C_9H_{16}N]^+$ | 0.0114 | | |
| VIP 55 | 3.1219 | 685.4383 | 1410 | $C_{37}H_{67}O_{10}P$ | $[M+H\text{-}H_2O]^+$ | 8.2 | 37.3 | 4 | PG(C31:4) ? | n.f. | 0.0086 | | |
| VIP 24 [c] | 3.0975 | 502.2896 | 864 | $C_{25}H_{44}NO_7P$ | $[M+H]^+$ | 6.3 | 32.2 | 2a | *Lyso* -PE(C20:4) | 361.2854 $[M\text{-}141.0189]^+$ $[M\text{-}C_2H_8NO_4P]^+$ $[C_{23}H_{37}O_3]^+$ | 0.0353 | | |
| VIP 56 | 3.0127 | 810.5935 | 1170 | $C_{46}H_{84}NO_8P$ | $[M+H]^+$ | 8.9 | 60.8 | 3 | PC(C38:4) | 184.0727 $[C_5H_{15}NO_4P]^+$ | 0.0469 | | |

Color code for normalized concentrations -1 1

### Supplementary figures

**Figure S1. Workflow of the complementary approaches, combining 16S and 28S rRNA gene barcoding of sponges, 16S rRNA gene metabarcoding of the microbiome, UHPLC-MS metabolomics of holometabolome.**

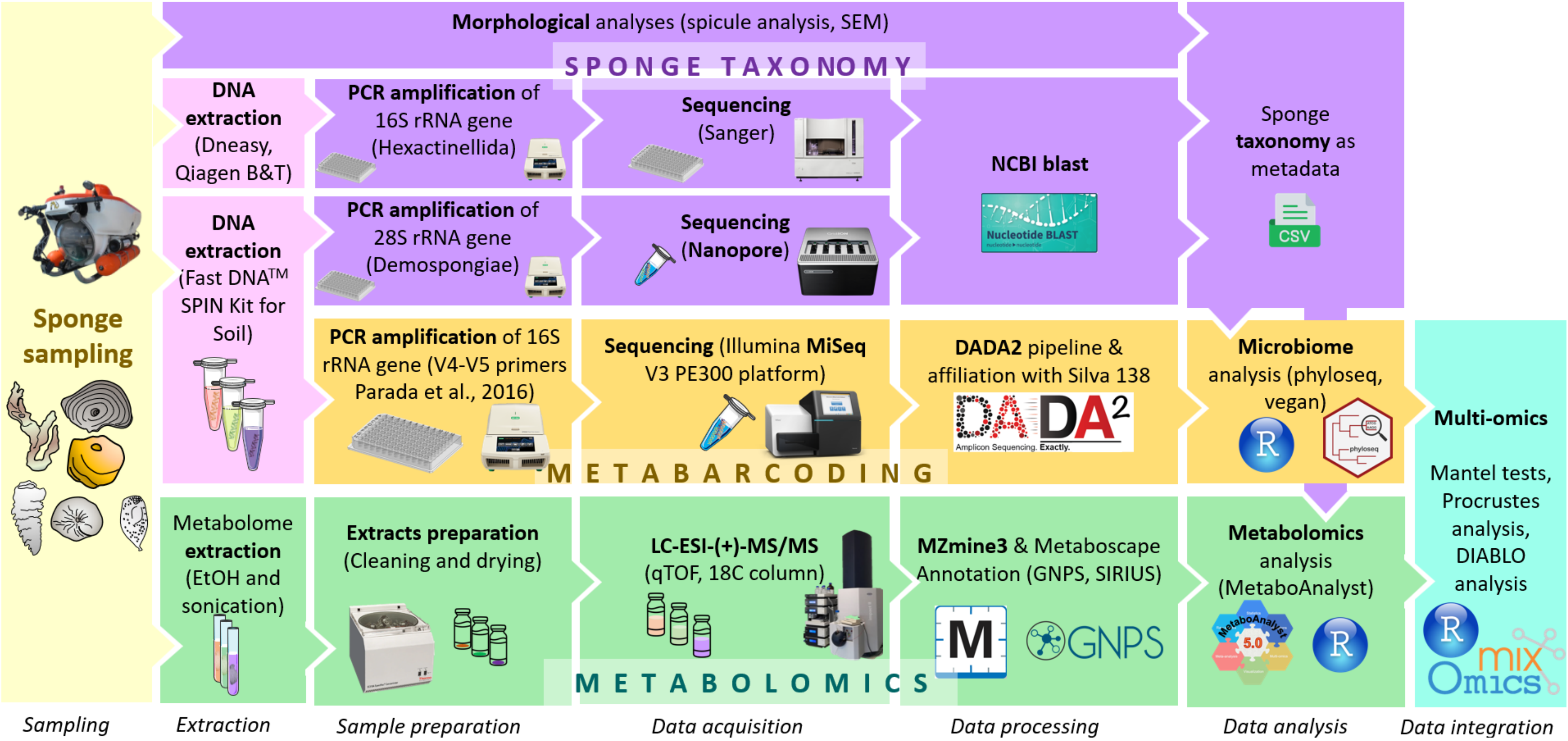

**Figure S2. Map of the sampling site. A. Location of Curaçao and bathymetry data. B. Detailed map of the study area.** The red circle indicates the Curaçao substation location. The red polygon indicates the reef area covered during the subdives (approximately 0.5 km$^2$).

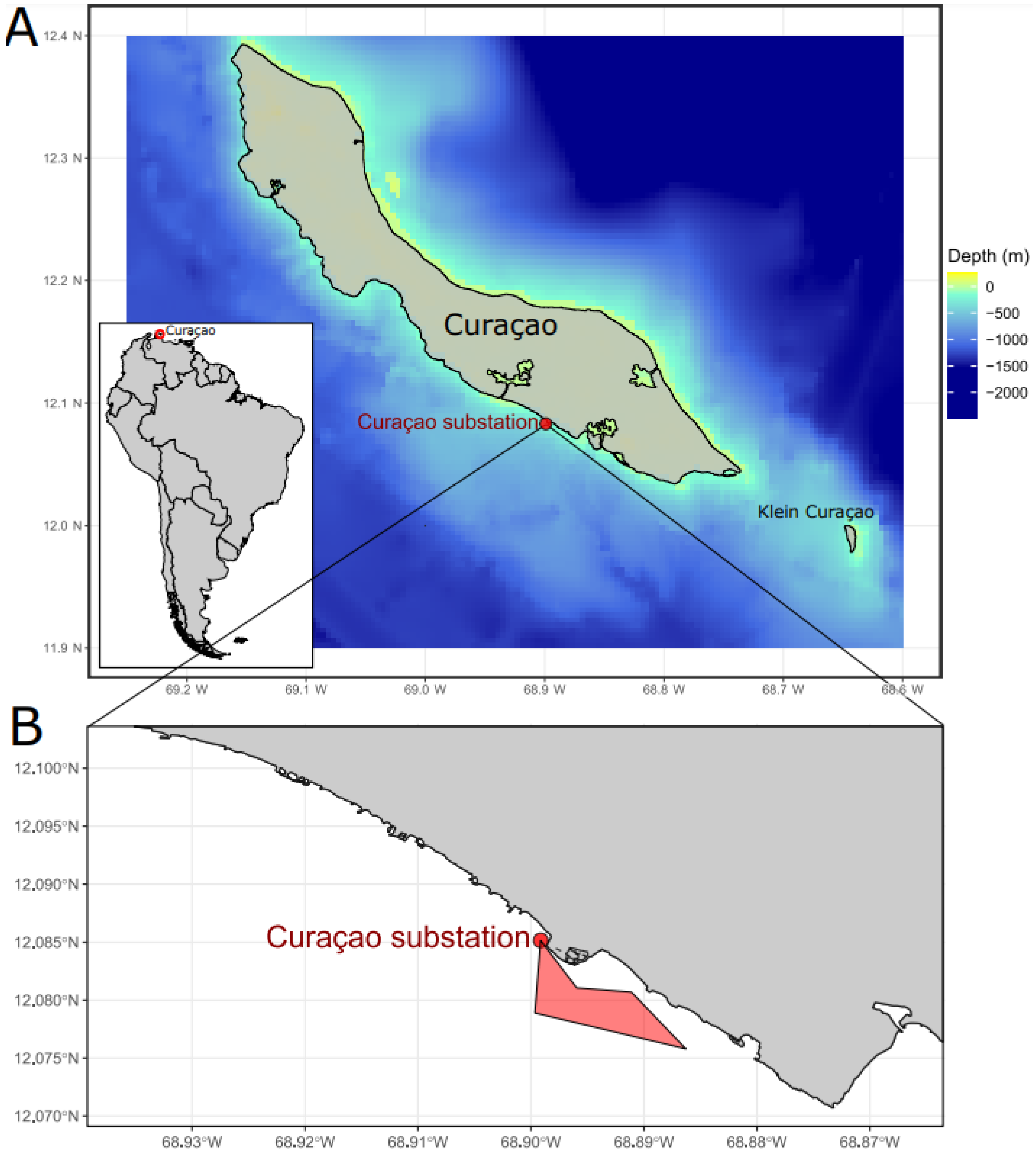

**Figure S3. Pictures of collected sponge specimens.**

**A.** Specimen of *Petrosia* aff. *weinbergi*. **B.** Specimen of *Neopetrosia eurystomata*. **C.** Frament of *Svenzea zeai* **D.** Specimen of *Cinachyrella kuekenthali* **E.** Specimen of *Geodia* aff. *Curacaoensis*. **F.** Specimen of *Petrosia* sp. ***G.*** Specimen of *Gastrophanella implexa*. **H.** Specimen of *Aciculites higginsii*. **I.** Specimen of *Topsentia ophiraphidites*. **J.** Fragment of *Calthropella lithistina*. **K.** Fragment of *Biemna microacanthosigma*. **L.** Specimen of *Geodia* cf. *megastrella*. **M.** Specimen of *Penares mastoideus*. **N.** Specimen of *Dactylocalyx pumiceus*. **O.** Specimen of *Leyfroyella sp*. **P.** Fragments of *Conorete pourtalesi*. **Q.** Specimen of *Verrucocoeloidea liberatorii*. **R.** Specimen of *Heterotella sp*. **S.** Specimen of *Myliusia callocyathus*. **T.** Specimen of *Hexactinella* sp. Scale bars on the bottom right sides of the pictures correspond to 2 cm.

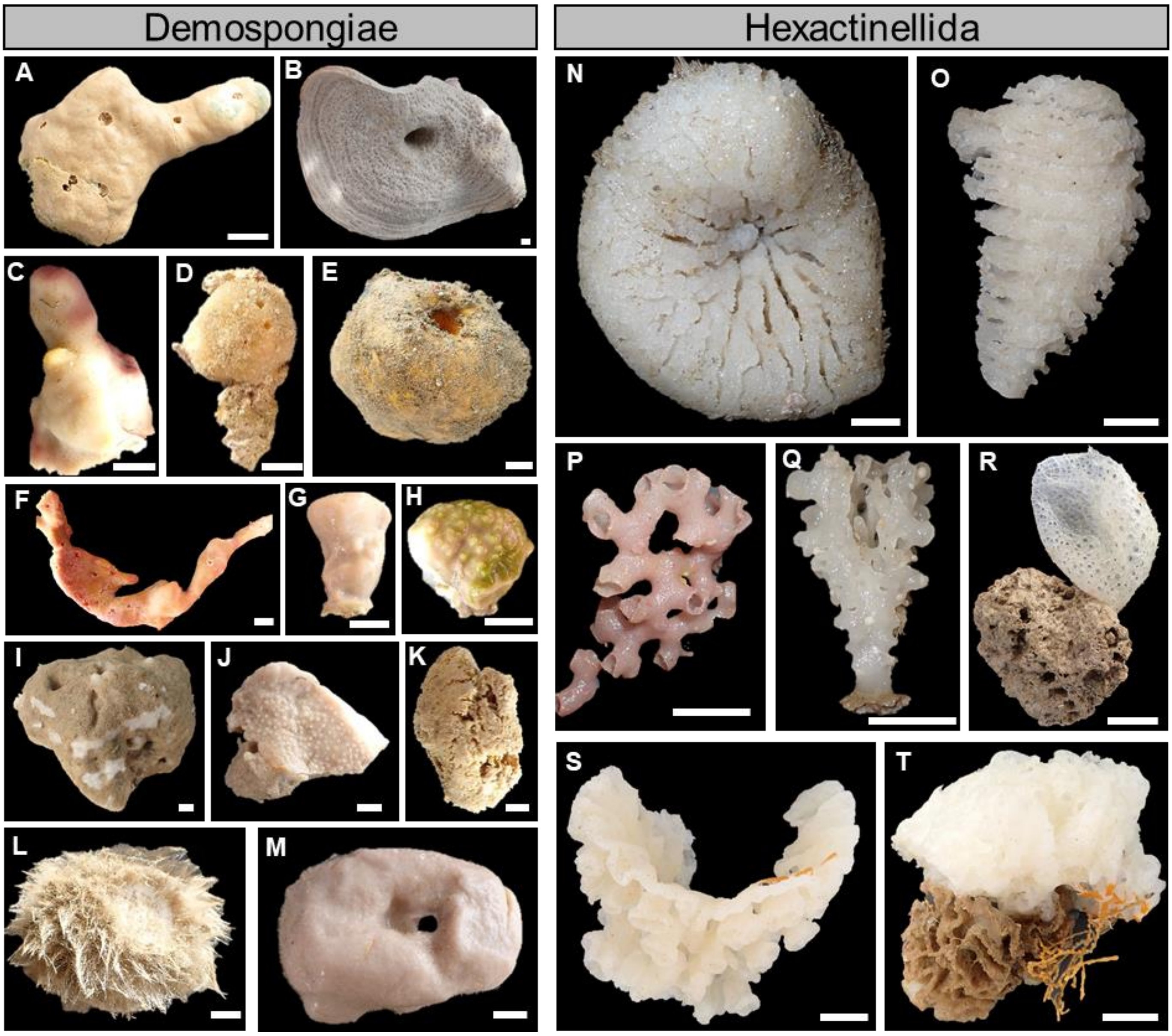

**Figure S4. Pictures of the hexactinellid-associated fauna.**

**A.** Picture of *in situ Verrucocoeloidea liberatorii* specimens colonized by the *Vitrumanthus schrieri* zoanthids. **B.** Picture of a collected *Verrucocoeloidea liberatorii* specimen colonized by the *Vitrumanthus schrieri* zoanthids. **C.** Pictures of a collected *Conorete pourtalesi* specimen colonized by the *Vitrumanthus schrieri* zoanthids. **D.** Picture of *Eiconaxius caribbaeus* specimens corresponding to a shrimp couple inhabiting the cavities of a *Conorete pourtalesi* specimen. **E.** Picture of a couple (male on the left, and female with eggs on the right) of *Eiconaxius caribbaeus* specimens. Red arrows indicate the location of the zoanthids. Blue arrows indicate the location of the shrimps. Scale bars on the bottom right sides of the pictures B, C, and E correspond to 1 cm.

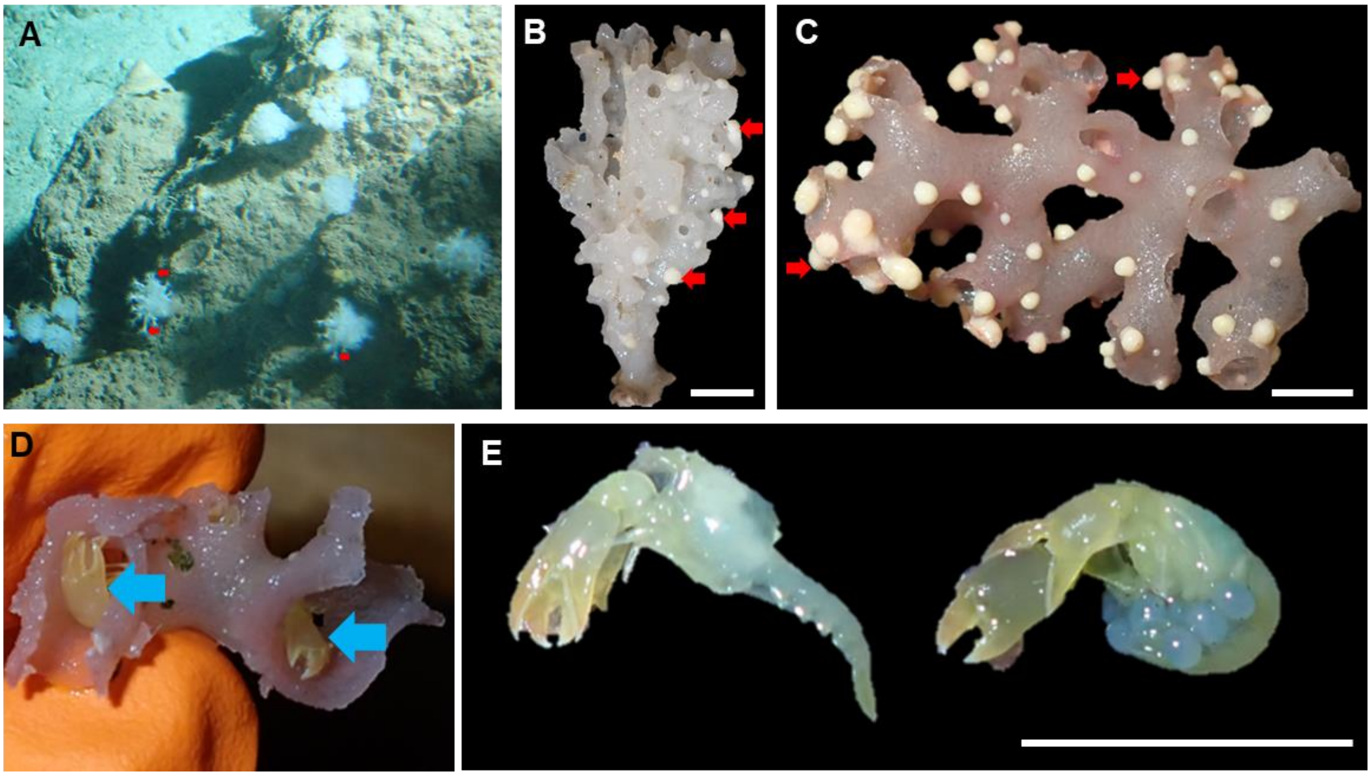

**Figure S5. $\alpha$-diversity analysis of the sponge-associated prokaryotic communities and holometabolomes.**

**A**. Rarefaction curves of the 16S rRNA gene reads. **B**. Boxplots of the $\alpha$-diversity metrics (Shannon, Chao1, and Pielou indices) of the prokaryotic communities associated with the Demospongiae and Hexactinellida samples in their respective photic zones (mesophotic, upper rariphotic, and lower rariphotic). Lowercase indices indicate results from the Wilcoxon pairwise test. **C**. Boxplots of the $\alpha$-chemodiversity measure (Shannon index) of the holometabolome associated with Demospongiae and Hexactinellida samples in their respective photic zones (mesophotic, upper rariphotic, and lower rariphotic). Lowercase indices indicate the results of Tukey's HSD pairwise test.

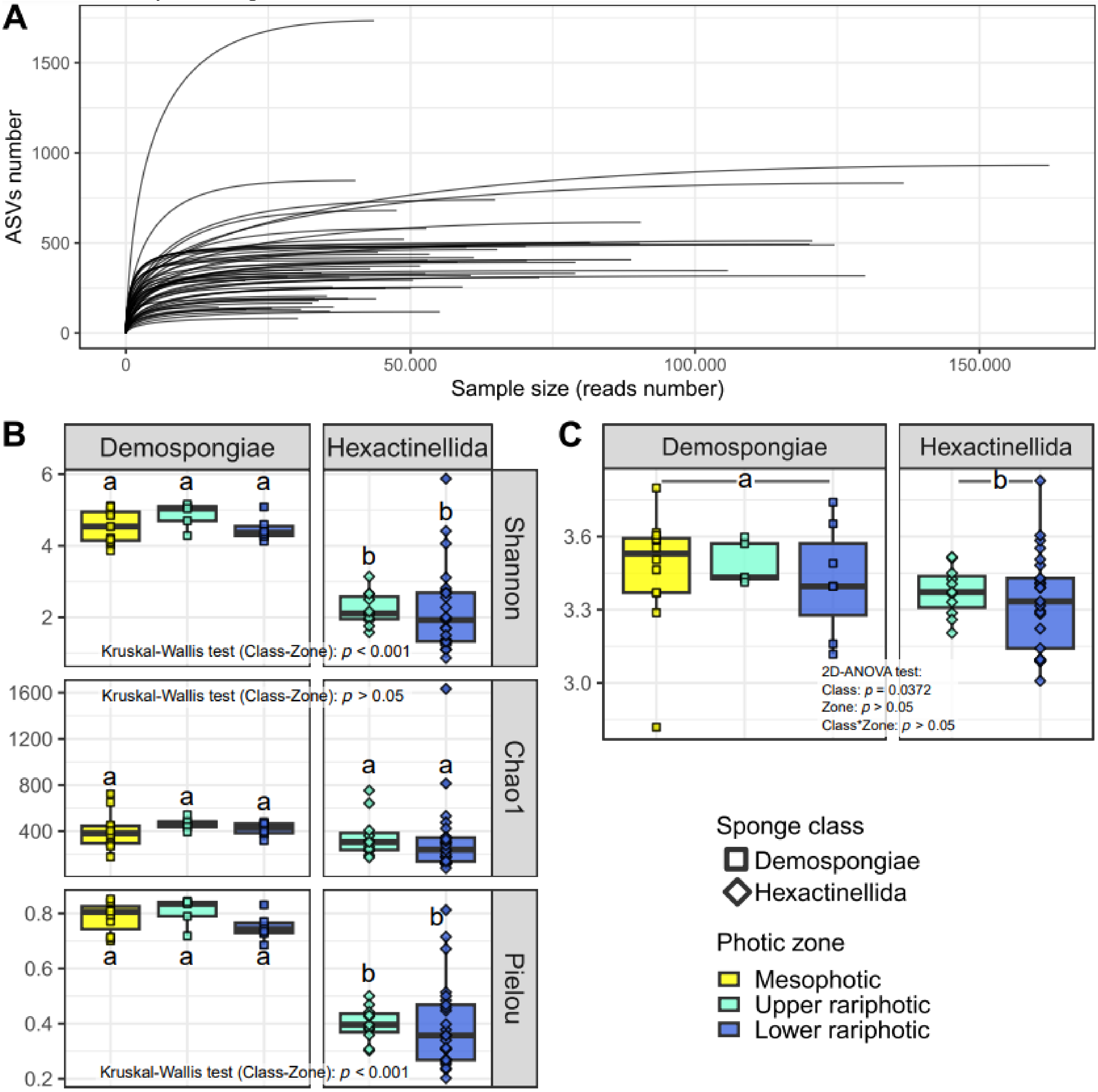

**Figure S6. Barplots of the prokaryotic community composition at the family level associated with the Demospongiae and Hexactinellida samples in their respective photic zones (mesophotic, upper rariphotic, and lower rariphotic).** "Other" corresponds to families below 5%. Sample name abbreviations are listed in Table S1

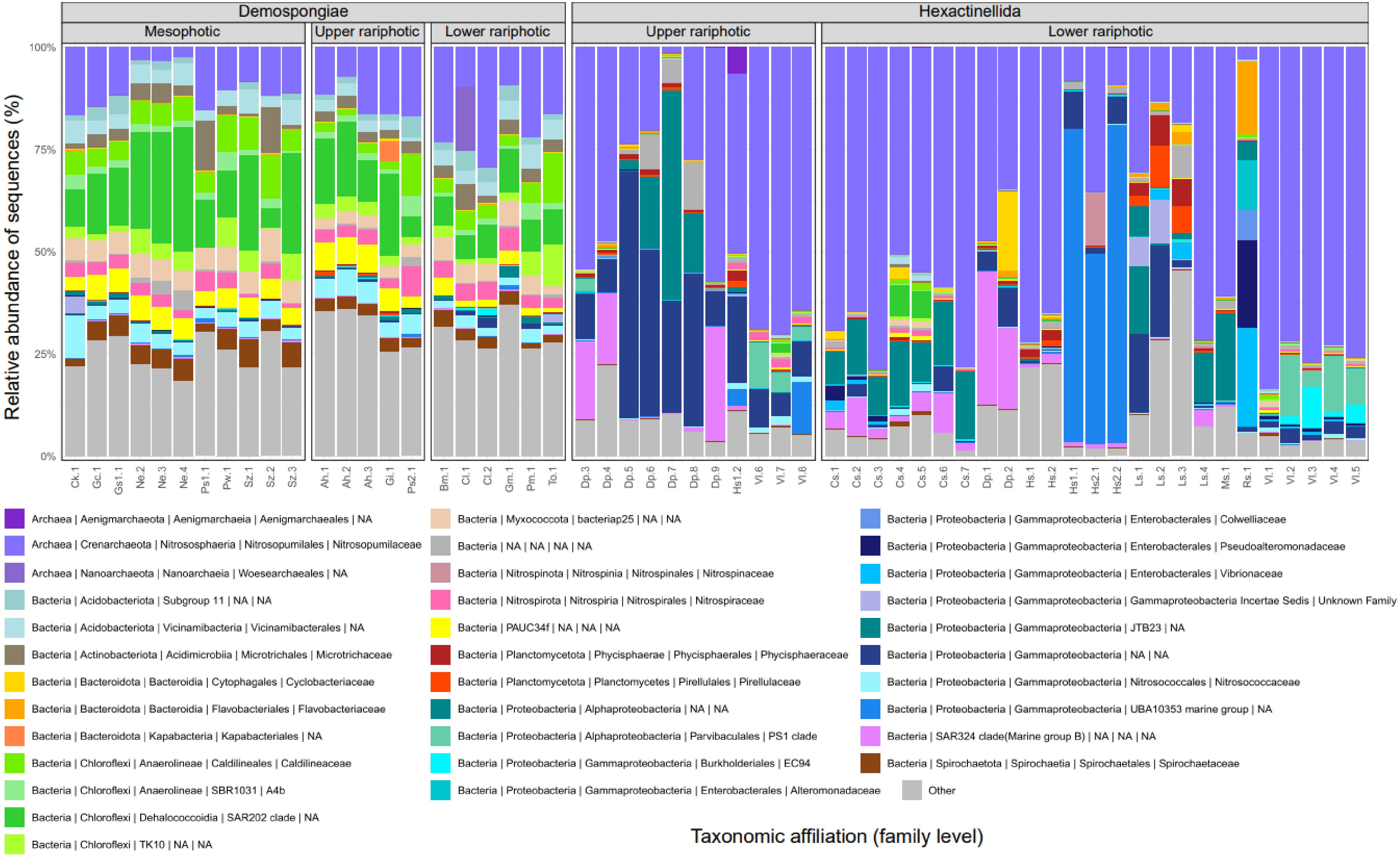

**Figure S7. Barplots of the archaeal community composition at the genus level associated with the Demospongiae and Hexactinellida samples in their respective photic zones (mesophotic, upper rariphotic, and lower rariphotic).** "Other" corresponds to families below 1%. Sample name abbreviations are listed in Table S1

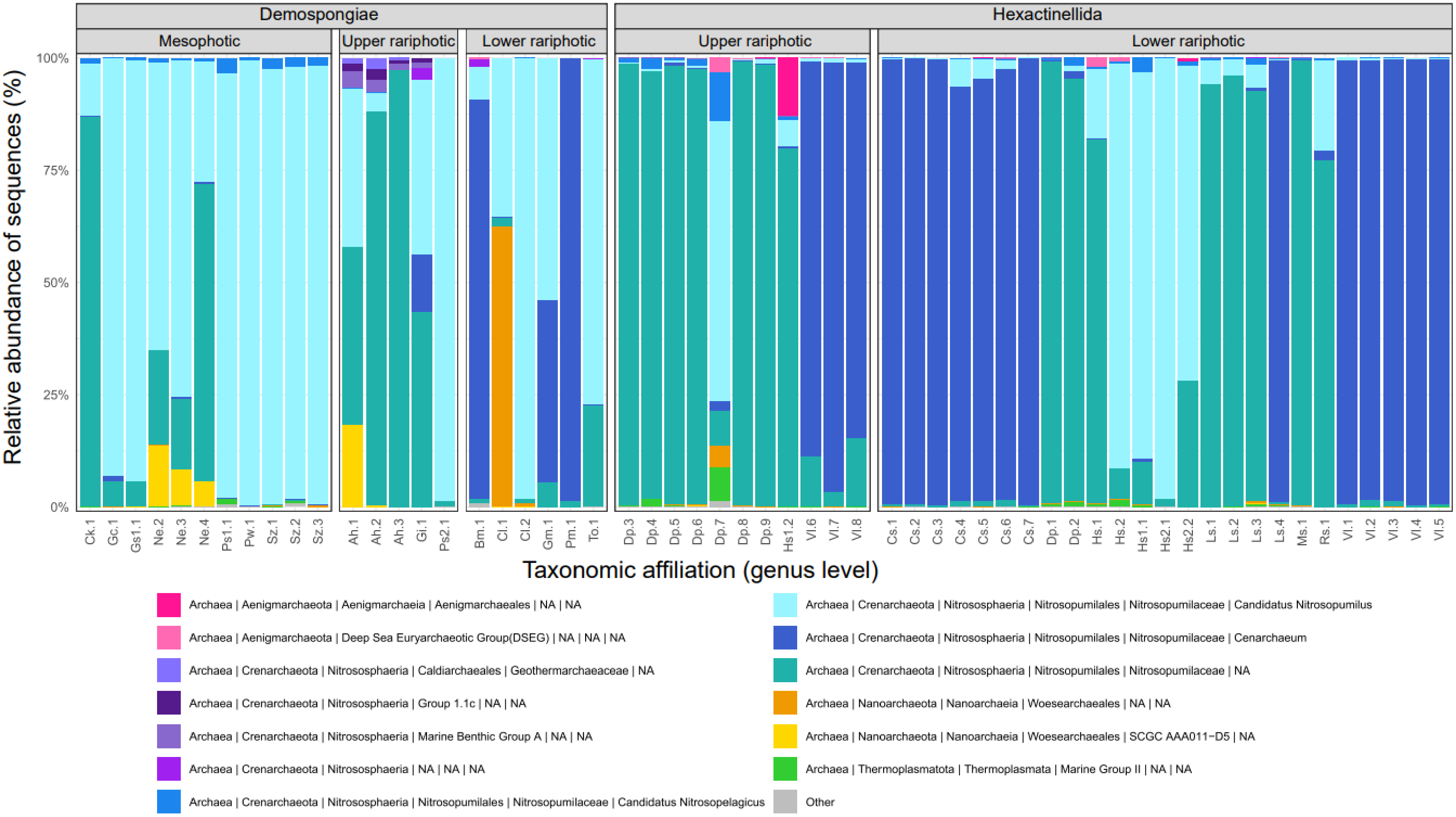

**Figure S8. Phylogenetical heat tree representing the taxa significantly and differentially abundant between Demospongiae and Hexactinellida samples.**

For each taxon, (i) the colors of their associated nodes correspond to the log2 fold change between Demospongiae and Hexactinellida samples, (ii) the size of the nodes corresponds to the relative abundance of each taxon.

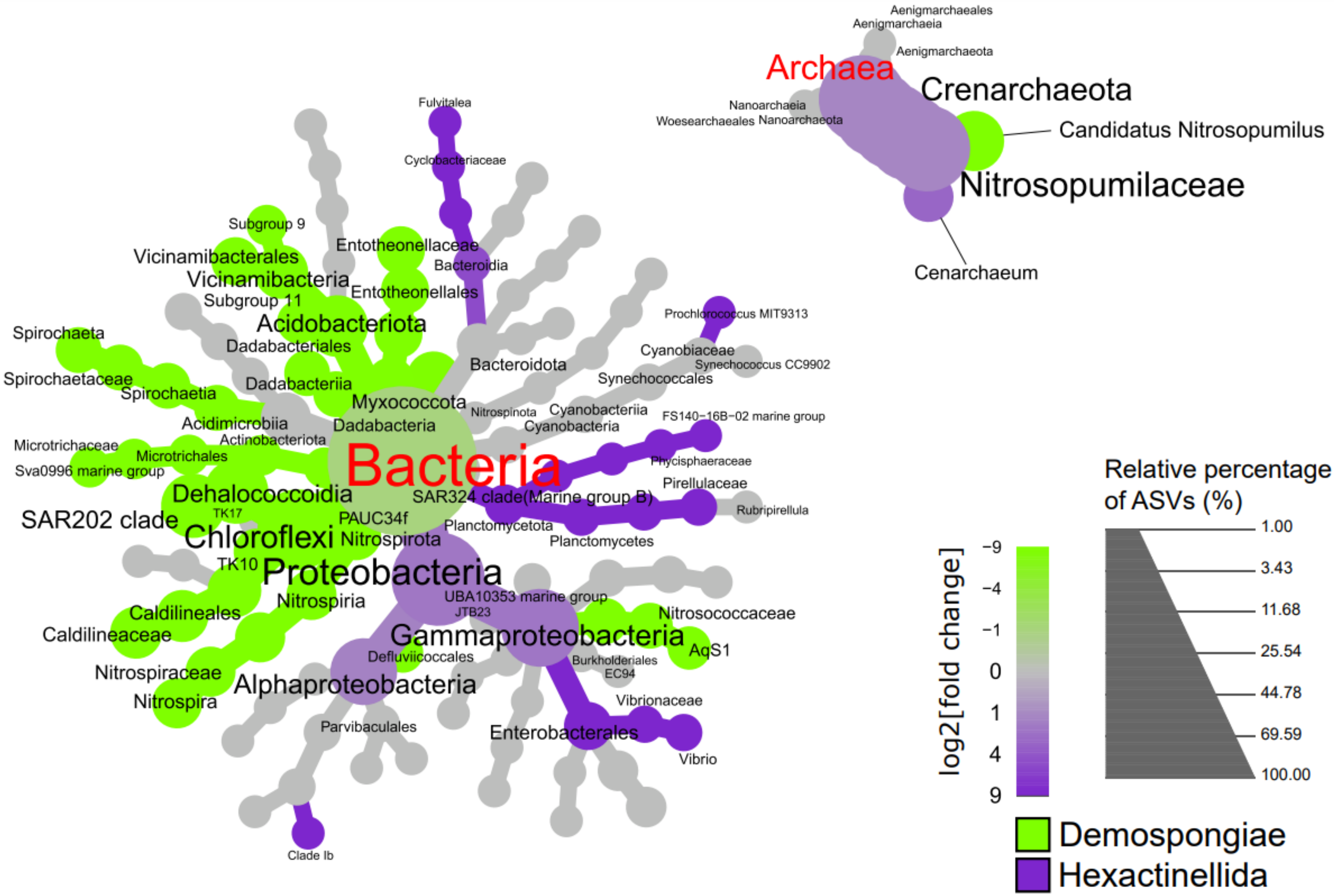

**Figure S9. Phylogenetical heat trees performed with Hexactinellida samples, and representing taxa significantly and differentially abundant between upper and lower rariphotic samples.**

For each taxon, (i) the colors of their associated nodes correspond to the log2 fold change between the ratio of the mean relative abundance within samples of each zone, (ii) the size of the nodes corresponds to the relative abundance of each taxon.

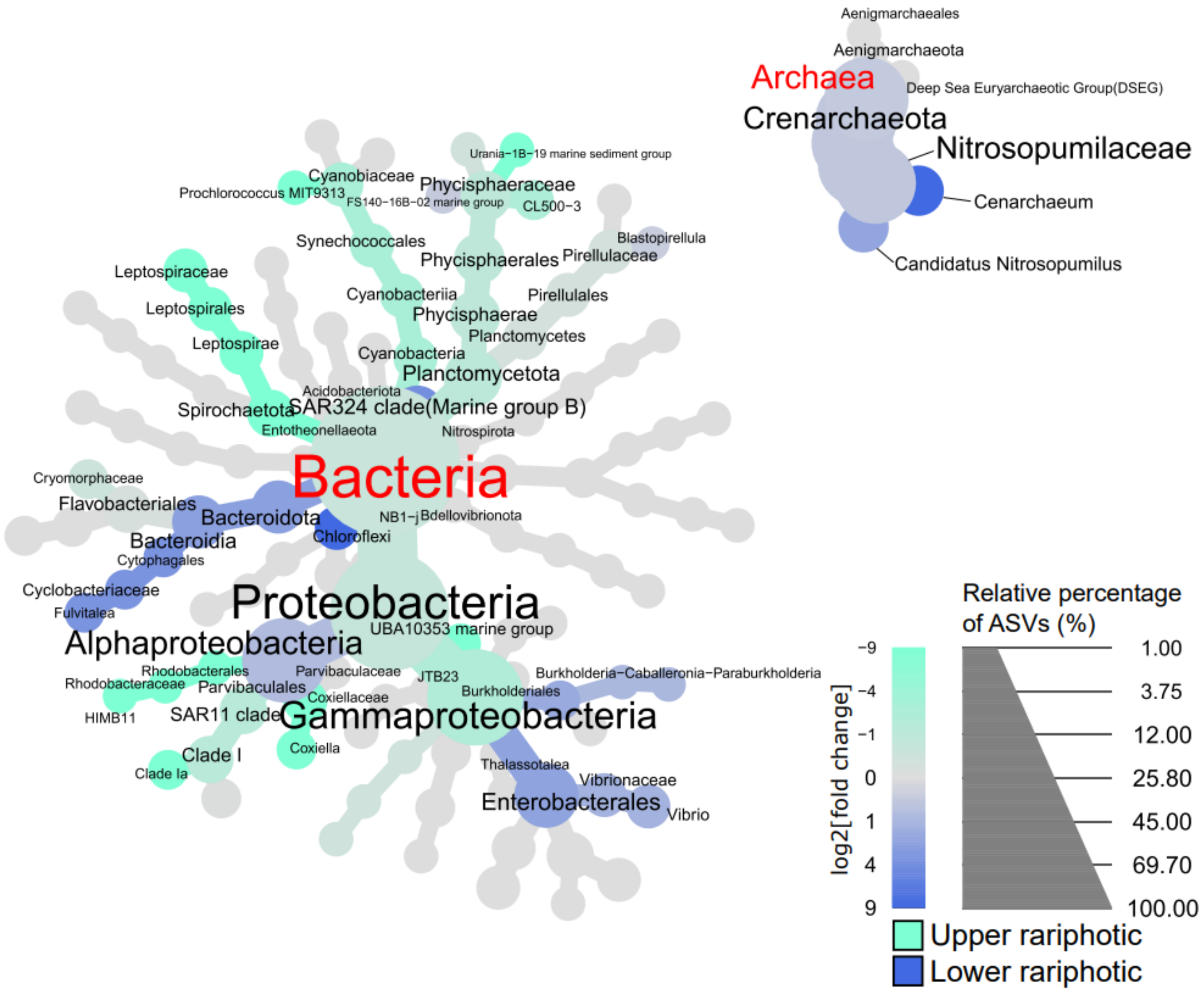

**Figure S10. PLS-DA plots of the holometabolomes conducted with the photic zone as a supervised factor, for both Demospongiae (A) and Hexactinellida (B) subseted datasets.**

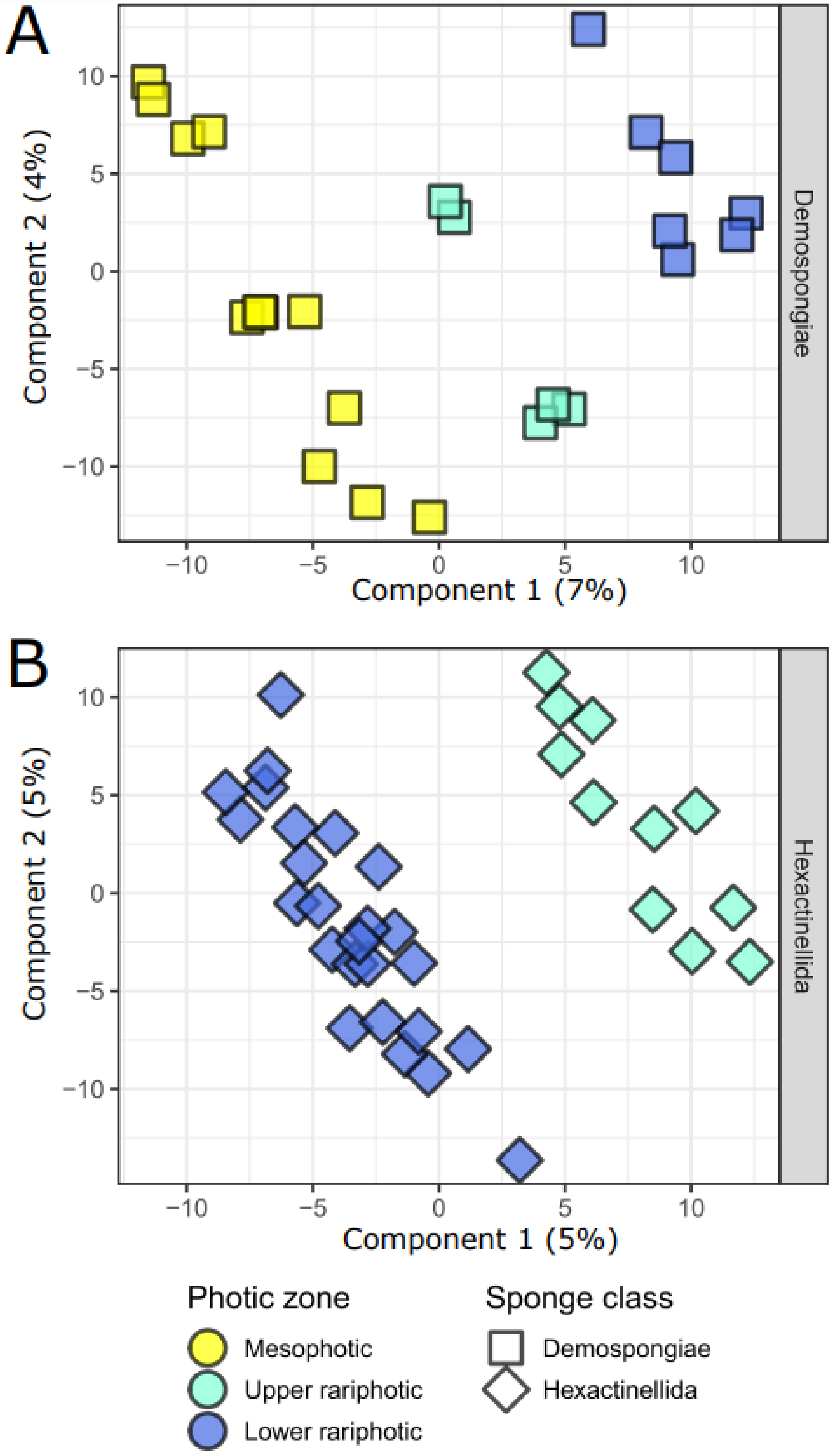

**Figure S11. FBMN analysis. A. Overall representation of the clusters (≥3 nodes) of the FBMN built with the MS/MS data.** Abbreviations: PC: phosphatidylcholine; PE: for phosphatidylethanolamine. The network also includes 32 clusters with 2 nodes and 1201 unbound nodes.

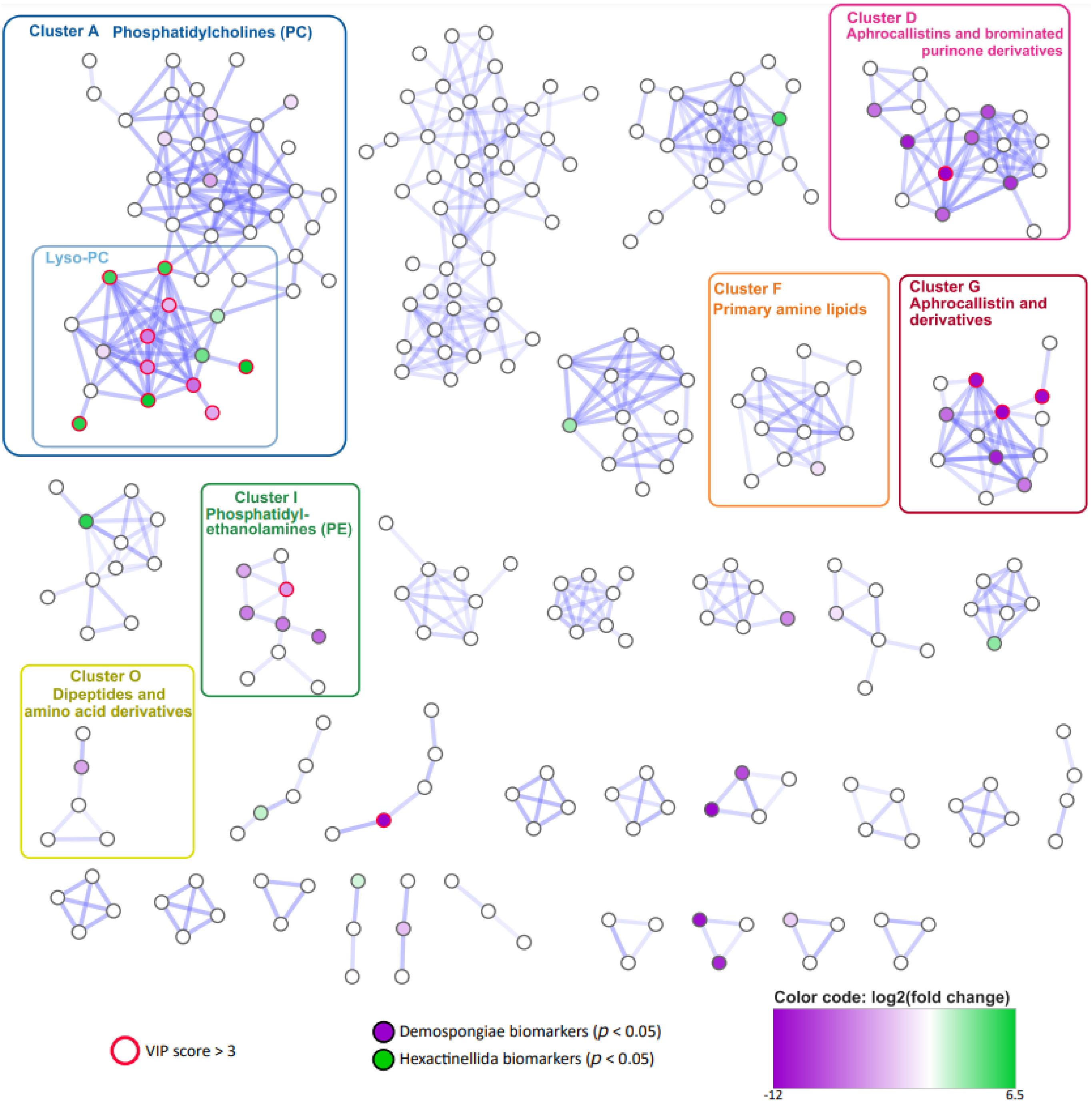

**Figure S12. Annotation of phospholipids identified within clusters A and D.** Abbreviations: PC: phosphatidylcholine; PE: for phosphatidylethanolamine.

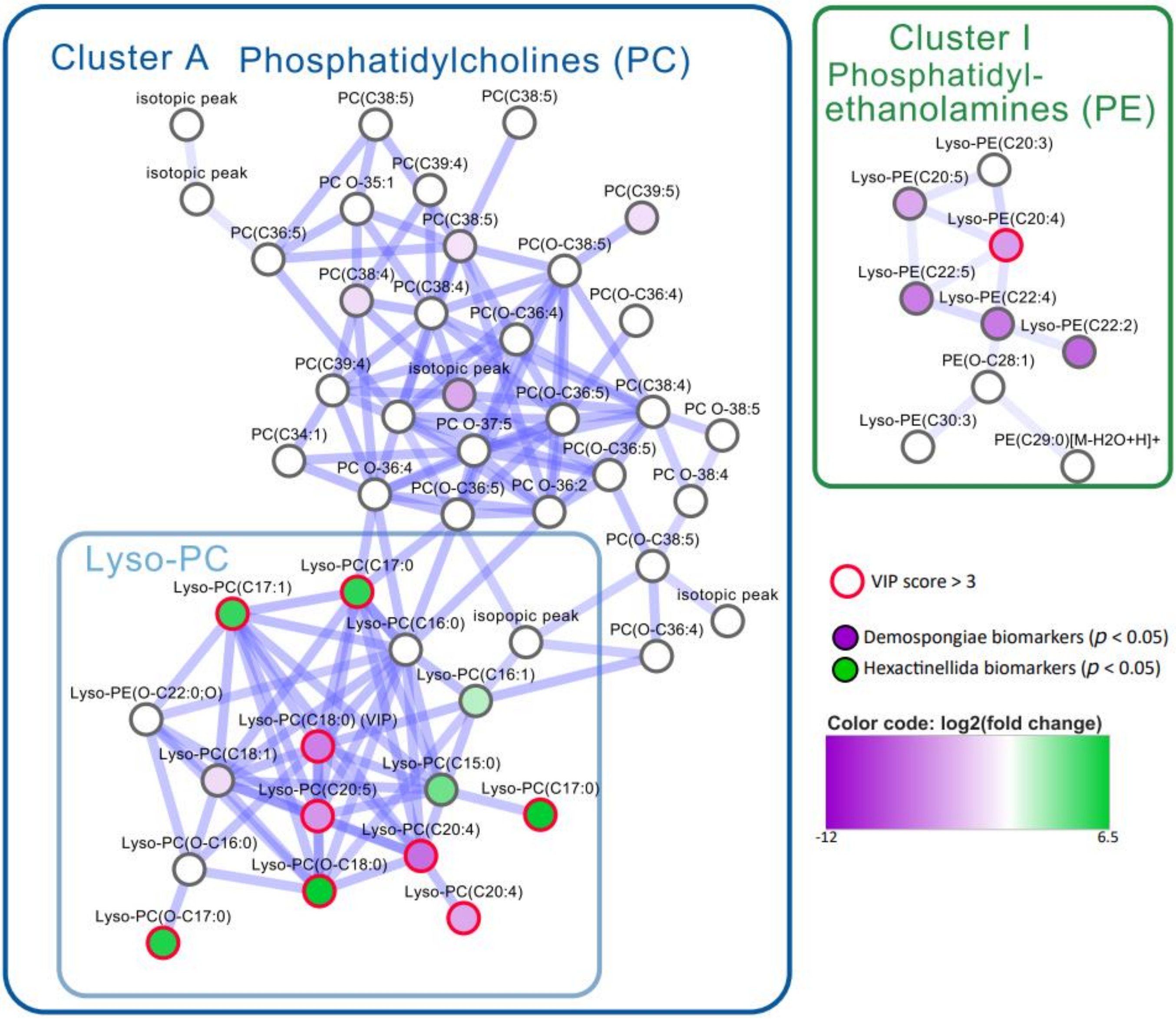

**Figure S13. Annotation of brominated compounds identified within clusters D and G.** The names "Aphrocallistin A", "Aphrocallistin B", and "Aphrocallistin C" were chosen according to Young et al., 2021. The name "Aphrocallistin" used in the manuscript and originally described by Wright et al., 2009 refers to "Aphrocallistin A".

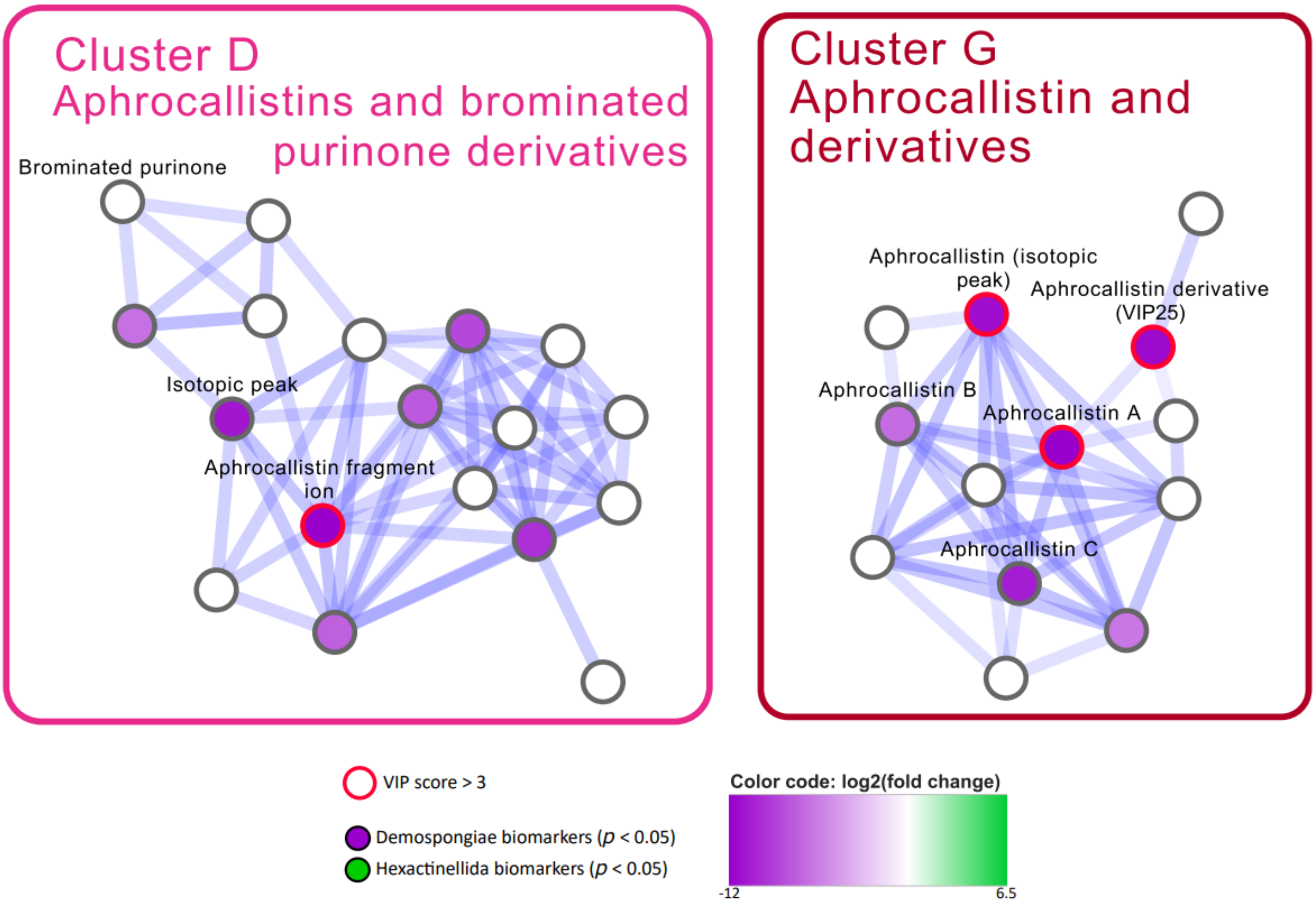

**Figure S14. HRMS/MS data (A) and putative MS fragmentation (B) of VIP n°13 identified as aphrocallistin (Wright et al., 2009).** The yellow box corresponds to a zoom on the 1:2:1 isotopic pattern of the precursor ion cluster, being characteristic of dibrominated compounds.

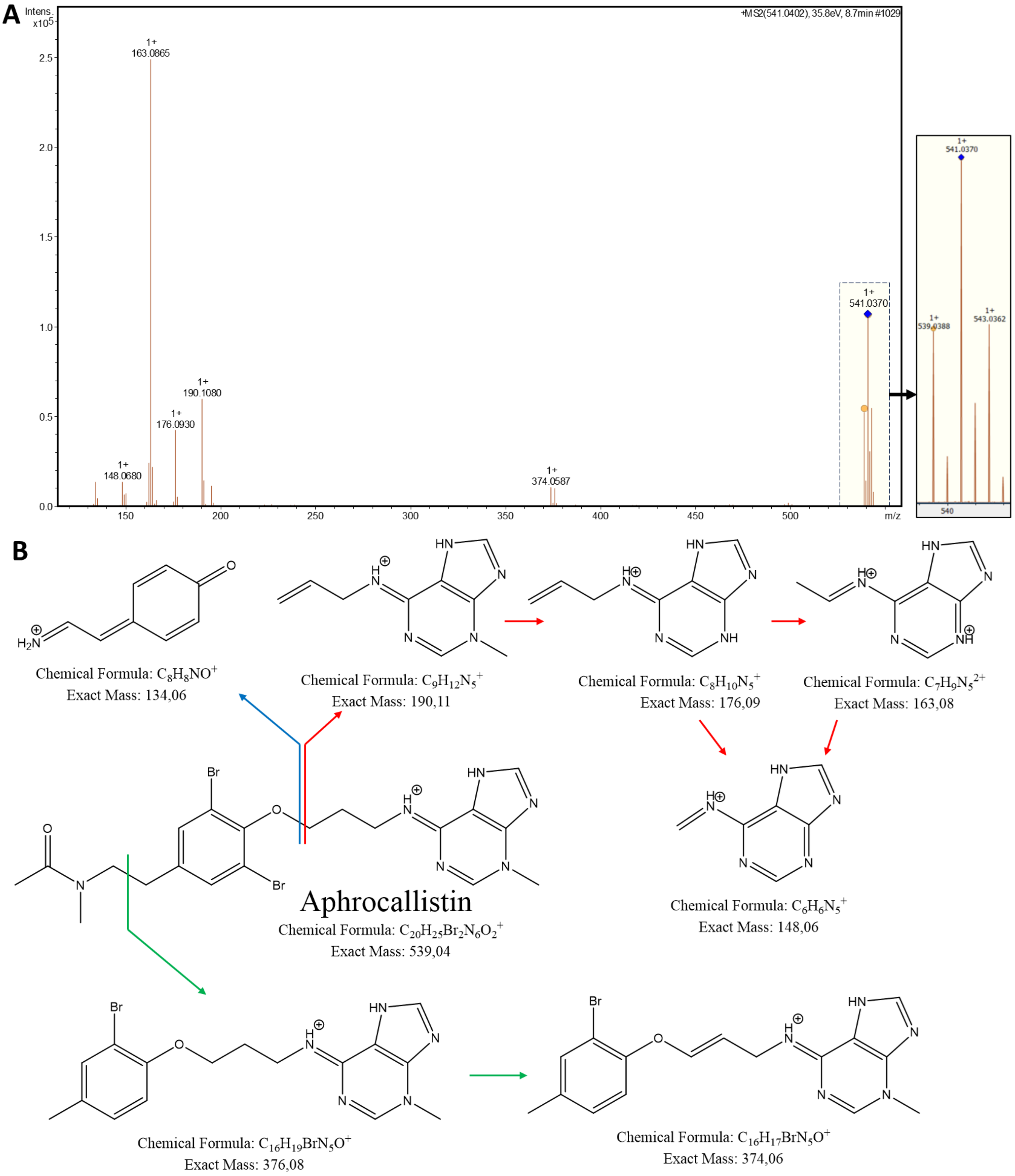

**Figure S15. Normalized concentrations of VIP n°13, 25, and 28, identified as aphrocallistin, an aphrocallistin derivative, and xanthurenic acid, respectively.** Lower case indices indicate the results of HSD Tukey's tests for pairwise comparison between genera.

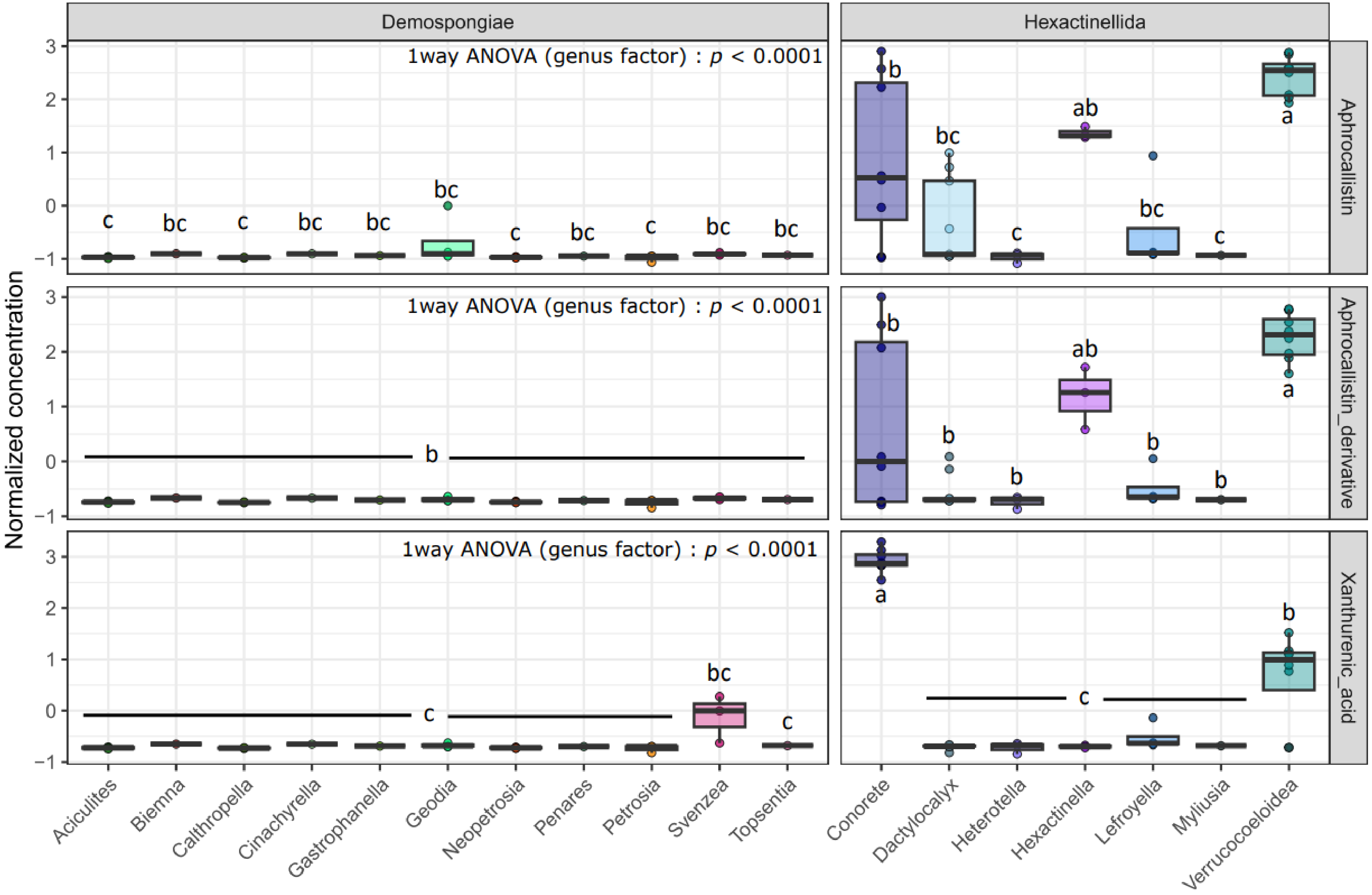

**Figure S16. HRMS/MS data and match with the GNPS Library Spectrum CCMSLIB of VIP n°27 identified as xanthurenic acid.**

This MS/MS data also matched the MS/MS data of xanthurenic acid within MassBank (MassBank Record: MSBNK-Fiocruz-FIO00659) and Bonnard et al., 2021.

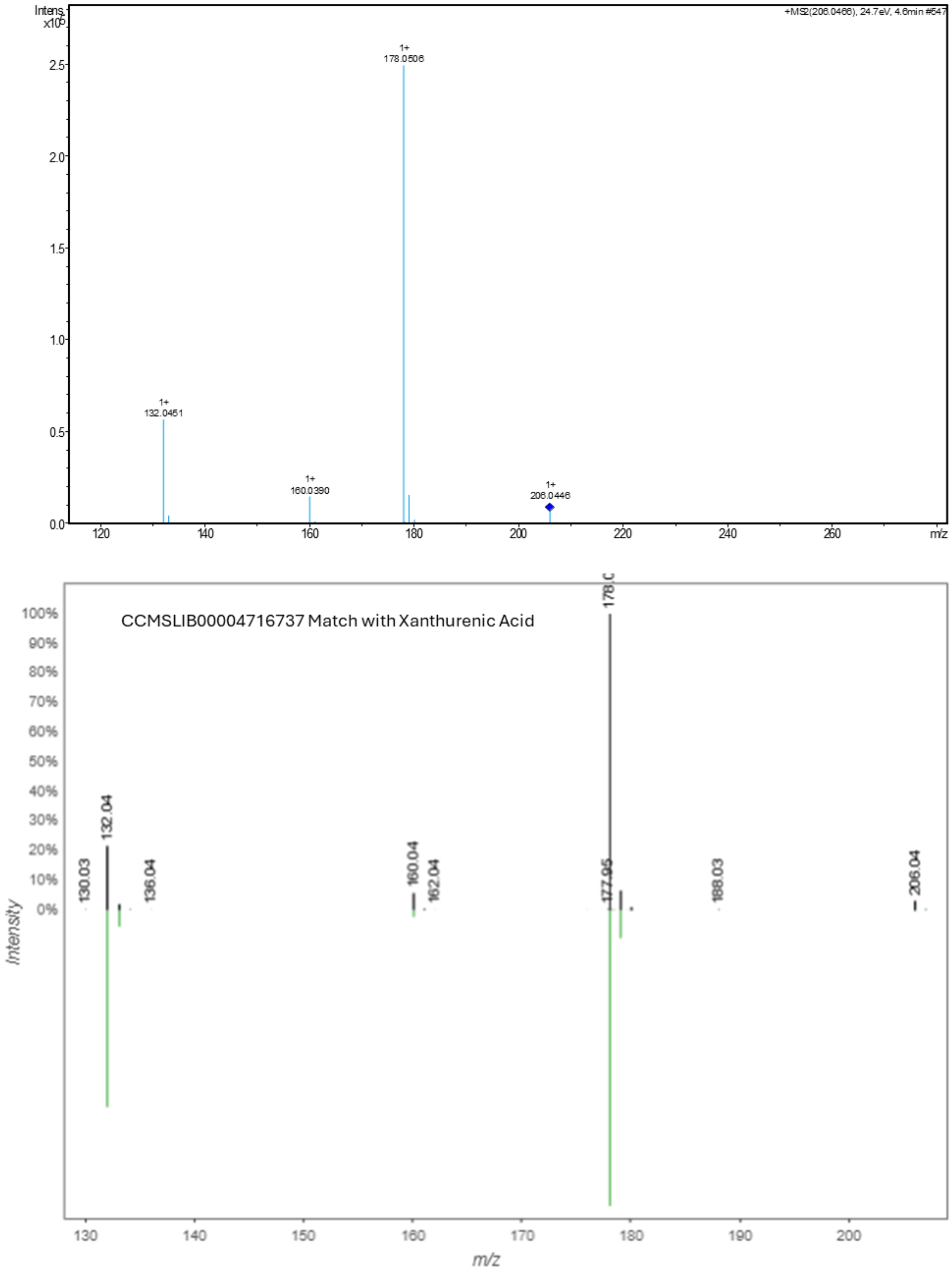
